# Enhanced transcriptional heterogeneity mediated by NF-κB super-enhancers

**DOI:** 10.1101/2021.07.13.452147

**Authors:** Johannes N. Wibisana, Takehiko Inaba, Hisaaki Shinohara, Noriko Yumoto, Tetsutaro Hayashi, Mana Umeda, Masashi Ebisawa, Itoshi Nikaido, Yasushi Sako, Mariko Okada

## Abstract

The transcription factor NF-κB, which plays an important role in cell fate determination, is involved in the activation of super-enhancers (SEs). However, the biological functions of the NF-κB SEs in gene control are not fully elucidated. We investigated the characteristics of NF-κB-mediated SE activity using fluorescence live-cell imaging of RelA, single-cell transcriptome, and chromatin accessibility analyses in anti-IgM-stimulated B cells. Cell stimulation induced nuclear foci formation of RelA and gene expression in a switch-like manner. The gained SEs induced a higher fold-change expression and enhanced cell-to-cell variability in transcriptional response. These properties were correlated with the number of gained cis-regulatory interactions, while switch-like gene induction was associated with the number of NF-κB binding sites in SE. Our study suggests that NF-κB SEs have unique roles for quantitative control of gene expression through direct binding to accessible DNA and enhanced DNA contacts.

## Introduction

The NF-κB signaling pathway, which plays important role in cellular functions and cell fate determination^1–3^, regulates proliferation and differentiation of B cells after B cell receptor (BCR) activation^4^. In BCR signaling, stimulation by antigens induces the canonical NF-κB pathway by activating protein kinase C β, kinase TAK1 (MAP3K7), BCL10, MALT1, and IκB kinase complex. This signaling cascade leads to the activation of IKKβ, which phosphorylates and promotes the proteasomal degradation of IκB. Since IκB masks the nuclear localization signal of NF-κB, its degradation promotes the nuclear translocation of p50 and RelA (p65) NF-κB heterodimer complex^5,6^. These proteins further act as transcription factors, promoting the transcription of NF-κB target genes.

Previous studies reported that anti-IgM stimulation induces activation of NF-κB pathway, eliciting an all-or-none response in the nuclear translocation of NF-κB at the single-cell level^7^ and causing cell-to-cell variability in transcriptional response^8^. This heterogeneity within the cell population may be responsible for the varying B cell phenotypes under the same environmental conditions^9,10^. Moreover, this heterogeneity in cell response upon NF-κB induction has also been observed in other cell types^11,12^, indicating the presence of common functions associated with the transcriptional regulation of NF-κB. Furthermore, a previous study reported that the anti-IgM-induced all-or-none NF-κB nuclear translocation response was observed to lead to the formation of nuclear aggregates^7^, hereby referred to as foci.

In this study, we hypothesized that these foci may be related to super-enhancer (SE)-mediated transcriptional regulation while aiming to explore the quantitative relationship between NF-κB foci and target gene expression. SEs are long clusters of enhancers, which have been reported to control cell identity and serve as nuclear platforms for the cooperative binding of transcription factors^13,14^. In previous chromatin immunoprecipitation sequencing (ChIP-seq) analyses using NF-κB and histone H3 lysine 27 acetylation (H3K27Ac) antibodies, NF-κB has been reported to be involved in SE activation along with other transcription factors^8,15^ and coactivators such as bromodomain protein 4 (BRD4)^16^. The aggregation of multiple transcriptional coactivators and mediators possessing intrinsically disordered regions (IDRs) promotes the formation of liquid-liquid phase separation (LLPS), which favors efficient gene transcription^17^. Together with RNA polymerase II, these proteins have been observed under fluorescence microscopy^17,18^ to form LLPS condensates at enhancer regions^19^ through the interaction of their activation domains^20^. However, there was no clear evidence that nuclear NF-κB aggregates present LLPS properties resembling SEs.

Anti-IgM-dependent NF-κB SE activity in primary splenic B cells caused higher fold change and threshold gene expression, inducing a wider distribution of gene expression in the cell population depending on the frequency of NF-κB binding to DNA^8^. NF-κB-mediated gene expression upon TNF stimulation has also been reported to present an enhanced gene expression heterogeneity through the modulation of transcriptional burst due to histone acetylation deposition and the presence of the initiation Ser5p RNA polymerase II^21,22^. In embryonic stem cells, it was reported that the elements causing noise in gene expression include promoter type, conflicting chromatin states, lack of active histone modification, and SE existence^23^. In contrast, a stochastic model of SE-mediated transcriptional regulation exhibit a low variation pattern in transcription noise than a typical enhancer (TE) does^24^. Therefore, it remains unclear whether SE can truly induce an enhanced transcriptional heterogeneity and if it is the case, what is the underlying mechanism resulting in heterogeneity.

In this study, we performed single-cell fluorescence imaging to confirm the visualization of NF-κB (RelA)-mediated SE formation and assessed the biochemical properties and dynamics of RelA foci upon anti-IgM stimulation in chicken DT40 B cells. We further utilized scATAC-seq (single-cell assay for transposase accessible chromatin with sequencing) to investigate the changes in SE activity upon cell activation through chromatin accessibility, and predicted the cis-regulatory interactions through co-accessibility analysis^25^. We observed that the formation of NF-κB nuclear aggregates upon nuclear translocation in anti-IgM-stimulated cells exhibits switch-like dynamics and was sensitive to a BET bromodomain inhibitor JQ1 and an LLPS inhibitor 1,6-hexanediol. Single-cell RNA and ATAC sequencing analyses further revealed that fold-change expression of the target genes and the enhanced transcriptional heterogeneity are correlated with the number of gained cis-regulatory interactions after cell stimulation. Our study also confirmed that BCR-mediated switch-like gene expression is associated with the number of putative NF-κB binding sites in a SE region. The overall results suggest that these slightly distinct quantitative transcriptional responses are originated from two mechanisms mediated by NF-κB: Direct binding to DNA and the biomolecular condensate assembly leading to enhanced DNA contacts. These findings clearly show that the biological functions of SE emerge from a combination of chromatin status and binding properties of transcription factors to facilitate divergent cellular responses.

## Results

### NF-κB proteins form SE-like nuclear foci

Initially, we observed under a fluorescence microscope the NF-κB foci formation in DT40 cells upon nuclear translocation of GFP-tagged RelA protein, a component of the NF-κB heterodimer. We observed that nuclear NF-κB maximally formed foci 20–30 min after anti-IgM stimulation (Fig. 1a, Supplementary Fig. 1a, b). As expected, an anti-IgM dose-dependent effect was observed in the numbers of RelA foci per cell, presenting a bimodal distribution across doses (Supplementary Fig. 1c), which was observed in a previous study^7^. Fitting the median number of foci of the dose-response curve to the Hill function resulted in a Hill coefficient of *N* = 4.33, suggesting that the formation of foci proceeded cooperatively (Fig. 1b). Since cooperative binding of transcription factors is a marker of SE-mediated gene regulation^17,26^, our results suggest that NF-κB may also activate the target genes cooperatively through these foci formation.

**Figure 1.**
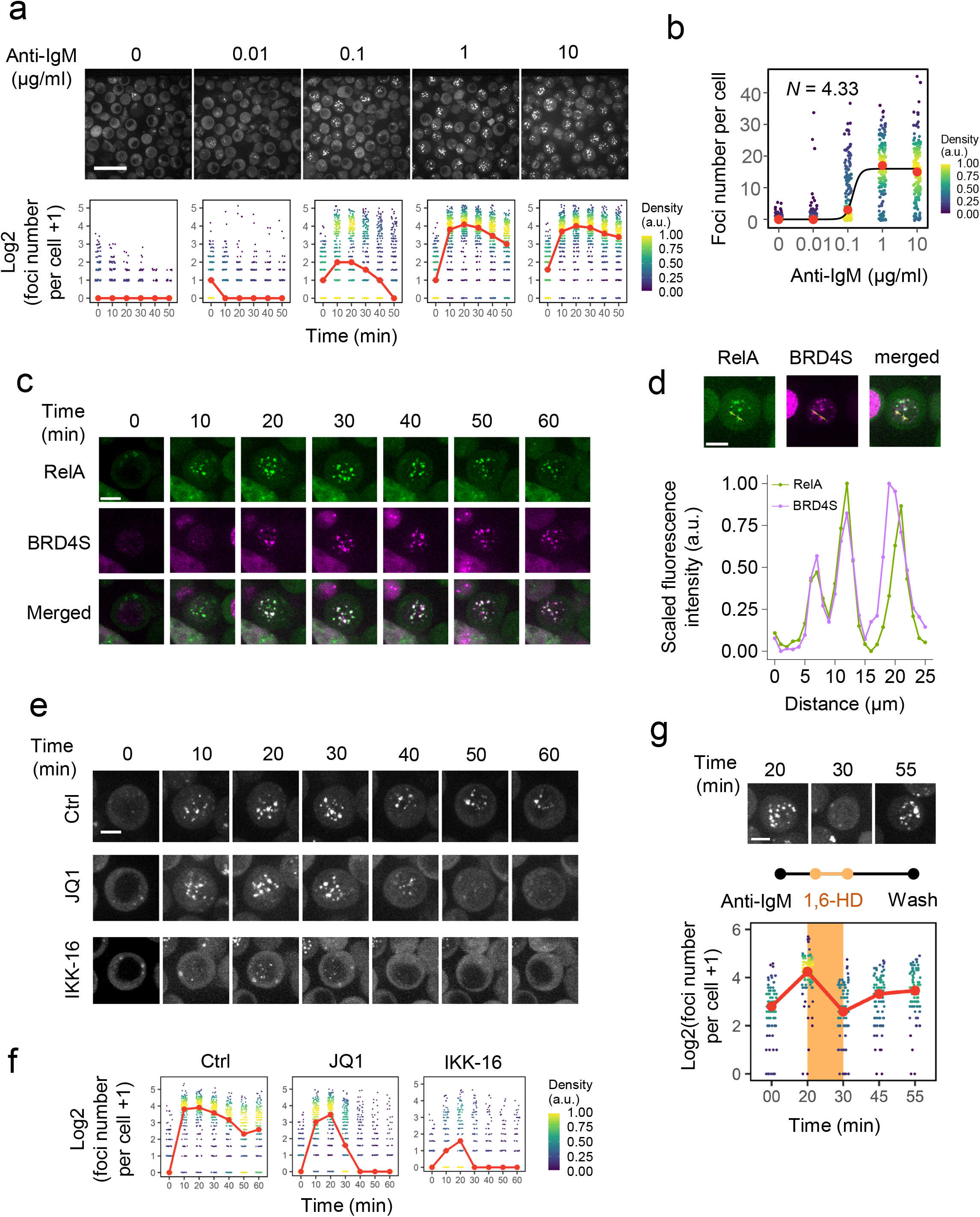
RelA foci demonstrate SE-like properties. (a) Representative real-time fluorescence micrographs of an individual RelA-GFP-expressing DT40 cell upon stimulation with 10 μg/mL anti-IgM (scale bar, 25 μm) and the quantification of the number of foci per cell across various anti-IgM doses (red points represent the median). (b) Quantification of the number of foci per cell after 20 min of various doses of anti-IgM stimulation (red dots represent the median). The median value was fitted to the Hill equation, resulting in a Hill coefficient of 4.33. (c) Time-lapse fluorescence micrograph of DT40 cells co-expressing mKate2-BRD4S and RelA-GFP upon stimulation with 10 μg/mL anti-IgM (scale bar, 5 μm). (d) Quantification of the co-localization of mKate2-BRD4S and RelA-GFP 20 min after anti-IgM stimulation. (e) Representative fluorescence micrographs of RelA-GFP-expressing DT40 cells pre-treated with 5 μM JQ1 for 60 min or 4 μM IKK-16 for 60 min. (f) Quantification of RelA foci from (e). (g) Quantification of RelA-GFP foci before treatment, after 1,6-hexanediol treatment and washing. The threshold of foci detection was lowered to compensate for the low fluorescence intensities of the recovering foci. (i) Density plot of *CD83* and *NFKBIA* single-cell expression obtained from scRNA-seq and smRNA-FISH after stimulation with 10 and 0 μg/mL anti-IgM.

To further investigate the properties of the RelA foci, we focused on the relationship between NF-κB and BRD4, which is known to bind to active SEs^27^. We generated DT40 cells co-expressing RelA-GFP and mKate2-BRD4S (see Methods). These cells revealed colocalization of BRD4S and NF-κB upon anti-IgM stimulation in a time-dependent manner (Fig. 1c–d). We further inhibited BRD4 activity by adding JQ1, a BET bromodomain inhibitor, 60 min before anti-IgM stimulation. The number of RelA foci increased to a maximum point at 20 min, where the difference between the control and JQ1 treated cells was negligible, then significantly decreased afterwards (Fig. 1e–f). At the same time, JQ1 inhibition disrupted BRD4S puncta formation (Supplementary Fig. 2). This indicates that RelA foci formation itself is BRD4-independent, but foci maintenance is BRD4-dependent. This is consistent with a previous result presenting that BRD4 maintained active NF-κB through RelA binding^16^. Conversely, inhibition of IκB kinase (IKK) using IKK-16 prevented the formation of RelA foci altogether (Fig. 1e–f), confirming that RelA foci formation is NF-κB pathway-dependent.

We further investigated the presence of LLPS-like properties in RelA foci. Earlier experimental approaches examined the presence of IDRs in RelA^28^. We used PONDR VLXT^29^ to analyze the IDRs and observed that RelA had high disorder scores in 250– 500 amino acid residues (54.1% of all residues), which was more or less comparable to BRD4 (85%) (Supplementary Fig. 3a). To examine whether the RelA foci exhibited LLPS-like properties, we treated cells with 1,6-hexanediol (1,6-HD), a compound known to promote the dissolution of liquid-like condensates^30^. Fluorescence imaging analysis (Fig. 1g, Supplementary Fig. 3b) presented that treatment with 1,6-HD after 20 min of anti-IgM stimulation dramatically reduced the number of foci and that washing the medium recovered the foci. These results demonstrate that the RelA foci have similar properties to SE and may be responsible for anti-IgM-dependent SE formation and target gene expression.

### Enhancement of heterogeneity in gene expression upon anti-IgM stimulation

To examine the above possibility, we next analyzed the transcriptional responses of DT40 single-cell populations (n=453) after one hour exposure to various anti-IgM doses (0, 0.01, 0.1, 1, and 10 μg/mL; Supplementary Table 1) using RamDA-seq, a method for single-cell RNA sequencing^31^ (scRNA-seq). Using Seurat, we obtained two distinct cell clusters (Fig. 2a)^32^. This suggests the presence of two discrete cell populations over the different doses of anti-IgM, such as those obtained in the imaging analysis (Supplementary Fig. 1c). We further performed a pseudo-time analysis to confirm that two cell populations were discrete (Supplementary Fig. 4a-b). We classified these clusters as activated (red cluster) and inactivated (blue cluster), as determined by their mean expression levels of representative NF-κB target genes, such as *NFKBIA, CD83*, and *TNFAIP3^33^* (Fig. 2b).

**Figure 2.**
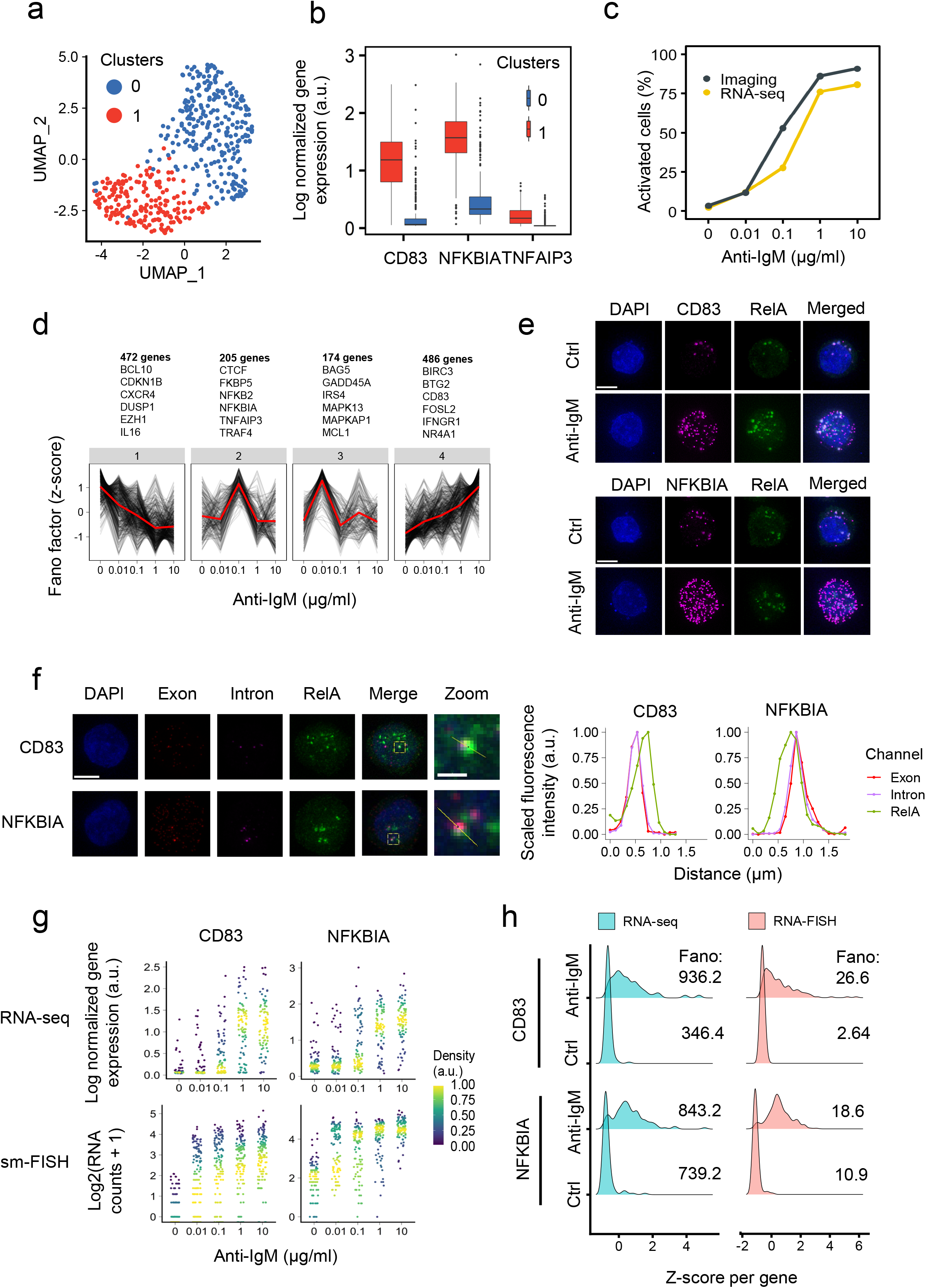
NF-κB SE-regulated genes demonstrate SE-like dynamics. (a) UMAP projection of the dimensionality reduction and clustering results of 453 cells scRNA-seq stimulated with various anti-IgM doses. (b) Box plot of the expression of known marker genes used to identify activated (red) and inactivated (blue) cell clusters. (c) Cell activation ratio from imaging obtained using logistic regression of foci at 20 min compared with the cell activation ratio from RNA-seq. (d) Hierarchical clustering analysis of Fano factors across anti-IgM concentrations for marker genes. The red lines represent the means. (e) Micrograph of *CD83* and *NFKBIA* smRNA-FISH (scale bar, 5 μm). (f) Micrograph of *CD83* and *NFKBIA* intronic smRNA-FISH (scale bar, 5 μm; zoomed scale bar, 1 μm). (g) RNA expression of *CD83* (B cell activation marker) and *NFKBIA* (NF-κB target gene) across dose points obtained from scRNA-seq (upper graph) and smRNA-FISH (lower graph). (h) Single-cell expression of *CD83* and *NFKBIA* obtained from scRNA-seq and smRNA-FISH after stimulation with 10 and 0 μg/mL anti-IgM.

Prediction of cell activation through logistic regression analysis on the number of foci per cell revealed similar quantitative profiles in cell activation predicted from RNA sequence data, in which most cells were activated at 1 and 10 μg/mL and inactivated at 0 and 0.01 μg/mL doses of anti-IgM (Fig. 2c, Supplementary Table 2). This suggests that the cell states determined from the scRNA-seq and imaging analysis were closely related.

Since the cells exhibited digital states, we investigated the change of gene expression variability across different concentrations of anti-IgM stimulation to confirm if gene expression is correlated with phenotypical differences. Thus, we calculated the Fano factor of 1,337 differentially expressed genes (DEGs) that were upregulated in the activated cells compared with inactivated cells from the scRNA-seq analysis at each dose point. The Fano factor (Ω^2^/μ), which is typically used to calculate changes in transcriptional bursting^34,35^, provides a measure of deviation from the Poisson distribution where Ω^2^/μ = 1. Further, we clustered the genes by heterogeneity dynamics by scaling Fano factor changes per gene to identify patterns of gene expression heterogeneity across concentrations (Fig. 2d). We observed that representative NF-κB negative modulators, such as *NFKBIA*^2^ and *TNFAIP3*^36^, present in cluster 2, have high heterogeneity at 0.1 μg/mL and thus, correlate with cellular heterogeneity which is supposed to be highest at 0.1 μg/mL (Fig. 2c). Interestingly, cluster 4 presents an increasing heterogeneity across anti-IgM concentrations, having the highest heterogeneity at 10 μg/mL, where most cells are phenotypically similar (Fig. 2c). This cluster was observed to contain B cells activation-related genes, such as *CD83* and *BIRC3*. The pseudo-time analysis also revealed that *CD83* has a higher gene expression heterogeneity upon cell activation than *NFKBIA* (Supplementary Fig. 4c). After further inspection, we observed that cluster 4 presents a significantly high Fano factor across anti-IgM concentrations than other clusters (Supplementary Fig. 5a). More importantly, genes in cluster 4 were enriched for immune cell activation related functions (Supplementary Fig. 5b). From these findings, we selected *NFKBIA* and *CD83* as representative genes for further analysis.

To investigate the relationship between the RelA foci numbers and target gene mRNA in the same cell upon anti-IgM stimulation, we quantified their cellular mRNA using smRNA-FISH (single-molecule RNA-FISH) (Fig. 2e). There was a positive correlation between the RelA foci number and mRNA expression of both genes at all dose points (*R*=0.49 for *NFKBIA, R=0.52* for *CD83)* (Supplementary Fig. 6a–b) and thus, NF-κB foci formation is an indication of the expression of these target genes. We also confirmed that the RelA foci numbers per cell across doses of the *CD83* and *NFKBIA* smRNA-FISH samples were similar (Supplementary Fig. 6c), suggesting that the technical differences between the samples were minimal. It should be noted that the correlation coefficient in this study represents only a rough estimate of the correlation as there was a difficulty in accurately counting the number of RelA foci due to the degradation during smRNA-FISH sample preparation and differences in the timing of gene expression and RelA translocation. Furthermore, to investigate the relationship between the RelA foci and transcriptional activity, we performed intronic smRNA-FISH to visualize nascent transcripts of *CD83* and *NFKBIA* and observed colocalization between *CD83* and *NFKBIA* nascent transcripts and RelA foci (Fig. 2f).

For these 2 genes, we visualized the gene expression across different doses of anti-IgM stimuli through both smRNA-FISH (Fig. 2g) and RNA-seq (Fig. 2h). For *CD83*, we see a lower basal gene expression at 0 μg/mL anti-IgM than *NFKBIA* and an increasingly wide gene expression distribution across anti-IgM concentration (Fig. 2g, Supplementary Fig. 6c). In contrast, *NFKBIA* exhibited a bimodal gene expression at 0.1 μg/mL, while there was a uniformly high gene expression across cells at 1 and 10 μg/mL. To quantify these differences, we further calculated the Fano factor change between cells at doses of 10 and 0 μg/mL, where the cells were predominantly activated and inactivated, respectively (Fig. 2h). We observed a significantly higher fold-change increase in the Fano factor associated with *CD83* (RNA-seq: 2.7, RNA-FISH: 10.1) compared to *NFKBIA* (RNA-seq: 1.1, RNA-FISH: 1.7). In addition, the bimodality of *NFKBIA* expression was well reproduced using smRNA-FISH especially at an intermediate anti-IgM concentration (0.1 and 0.01 μg/mL), while *CD83* expression presented a single broad peak across different concentrations (Supplementary Fig. 7). This evidence indicates that the transcriptional regulation between these two genes upon cell activation is potentially controlled by different modes.

### Gained SEs are responsible for the strong induction of B cell development-related genes

To clarify the underlying transcriptional mechanism, we explored epigenetic regulation of NF-κB mediated SEs and their effects on transcription. Typically, SE is identified using rank-ordering of super-enhancers (ROSE) algorithm^13^ using transcription factors, H3K27Ac ChIP-seq, or DNase-seq data^13,14,26^. In our research, we utilized ATAC-seq data to investigate a large stretch of enhancers similar to SEs since both DNase-seq and ATAC-seq fundamentally identify open chromatin^37^ and the fold-change in H3K27Ac and ATAC signals are correlated in B cell SEs^8^.

Stitching of ATAC peaks was performed for peaks within 5 kb of each other, shorter than the default setting (12.5 kb) since ATAC peaks are often broader than TF ChIP-seq peaks^38^. The ROSE algorithm was further implemented^13,39^ to rank the enhancers of the cells with and without anti-IgM stimulation (Fig. 3a). Changes in chromatin accessibility between the unstimulated and stimulated cells were acquired by obtaining the merged peaks and calculating the fold changes of the signals between both conditions (see Methods). We observed that anti-IgM stimulation evoked a significant change in SE, but not in TE (Fig. 3b). Changes in SE signals can be observed in representative genes *CD83* and *NFKBIA* (Fig. 3a, c), and both the genes presented one of the highest gene expression fold changes among SE controlled genes (Fig. 3d). The ATAC signal fold-change is more correlated with gene expression fold-change in SE than TE (*R*=0.29 for SE, *R*=0.084 for TE) (Fig 3e). We confirmed that the SEs detected using ATAC-seq were comparable to the previous report using H3K27Ac ChIP-seq of mouse primary B cells^8^, where SEs have a longer genomic length than TEs (Supplementary Fig. 8a), and the length is highly correlated with the ATAC signal *(R* = 0.95) (Supplementary Fig. 8b). Further, the ranked SE associated genes, including *CD83* and *NFKBIA*, revealed a large fold-change RNA expression (Supplementary Fig. 9a) and increase in ATAC signal upon anti-IgM stimulation (Supplementary Fig. 9b) as identified from mouse primary B cell data, where the SEs were identified using H3K27Ac ChIP-seq data^8^. Therefore, we concluded that the SEs identified from ATAC-seq are comparable to the SEs identified from H3K27Ac ChIP-seq data in this experimental setting. Interestingly, *NFKBIA* was classified as a SE target in control (Fig. 3a) and presented fewer accessibility differences upon stimulation (Fig. 3c, e), indicating that *NFKBIA* expression might be less influenced by SEs and chromatin status than *CD83*.

**Figure 3.**
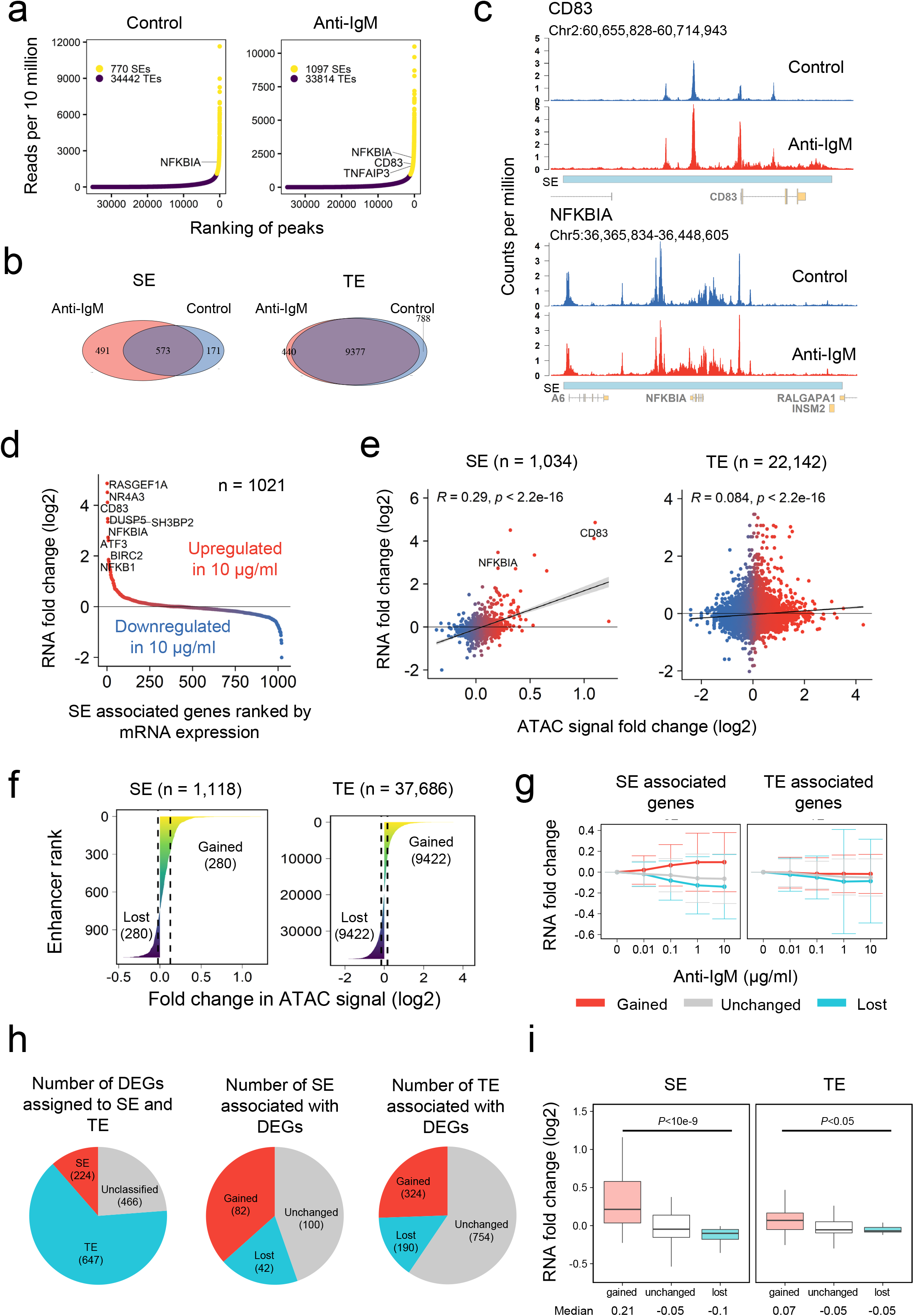
SE analysis. (a) Plot of SEs determined using the ROSE algorithm13 from cells with and without anti-IgM stimulation. (b) Venn diagram showing the number of SEs and TEs obtained from each stimulatory condition. (c) ATAC-seq track view of NF-κB target genes in B cell CD83 and NFKBIA SEs in control and 10 μg/mL anti-IgM stimulated cells. (d) Scatter plot of the mean fold-change of SE-associated genes between 10 μg/mL anti-IgM stimulated and control cells. (e) Correlation plot between TE and SE annotated peaks and the associated genes. Peaks associated with both SE and TE were assigned to SE. Genes with gene expression 0 across all doses were removed. (f) Regions with ATAC log2 fold changes signal more than the upper quantile were annotated as gained SEs/TEs and lower than the lower quantile were annotated as lost SEs/TEs. (g) Anti-IgM dose-response of fold change in the RNA level. Fold change was calculated from scRNA-seq by averaging the expression across all cells in the same stimulatory condition compared to dose 0. Number of genes: Gained SE, 260; Unchanged SE, 522; Lost SE, 239; Gained TE, 3598; Unchanged TE, 5384; Lost TE, 520. (h) Pie chart showing the classification of DEGs with SE and TE. (i) Boxplot of RNA log2 fold change between activated cells and inactivated cells for DEGs associated with TE and SE. Number of genes: Gained SE, 82; Unchanged SE, 100; Lost SE, 42; Gained TE, 242; Unchanged TE, 384; Lost TE, 21. *P*-values were calculated using one-way ANOVA with undersampling (n = 21).

We further classified the enhancers as gained, lost, or unchanged by the quantiles for TEs and SEs (Fig. 3f). In SE, mean RNA fold change is greater in higher dosage in gained SE and lost SE, while in TE, the difference is negligible (Fig. 3g). This result confirms that large chromatin structure changes are associated with dynamic gene regulation in SEs.

To investigate the role of SE in the whole transcriptome, we took the intersection of DEGs and SE and TE (Fig. 3h). Among all DEGs, 16.7% and 48.4% of DEGs are controlled by SE and TE, respectively, and about half of SE and TE are involved in the regulation of DEGs. As expected, gain and loss of SEs presented a higher upregulation and downregulation of associated DEGs, respectively, compared to those of TEs (Fig. 3i). Further, examination of biological functions of the gained SE-associated genes that have enhanced expression exhibited the enrichment of functions related to immune cell activation and regulation (Supplementary Fig. 10), confirming that SE-activated genes are responsible for B cell activation.

### SEs are also defined by threshold gene expression tied to NF-κB binding

NF-κB and PU.1 coexistence have been suggested to be one of the defining features of SEs in B cell activation^8^. In this study, we demonstrated that there are higher NF-κB (median motif occurrences: 16 for SE, 0 for TE; median motif occurrences per 10^4^ bp: 3.4 for SE, 0.0 for TE) and PU.1 (median motif occurrences: 50 for SE, 2 for TE; median motif occurrences per 10^4^ bp: 10.9 for SE, 8.1 for TE) motif occurrences at SE compared to at TE (Supplementary Fig. 11). Interestingly, between the enhancer categorization to gained, unchanged and lost, we observed that there is a difference in NF-κB motif density (4.7 for gained, 3.2 for unchanged, 2.7 for lost), but not in PU.1 (10.7 for gained, 10.8 for unchanged, 11.3 for lost) at SE (Fig. 4a-b). Therefore, this result updates the previous study^8^ where co-localization of NF-κB and PU.1 are relevant to SEs, albeit NF-κB has a stronger effect in determining changes in chromatin accessibility and SE formation after B cell activation.

**Figure 4.**
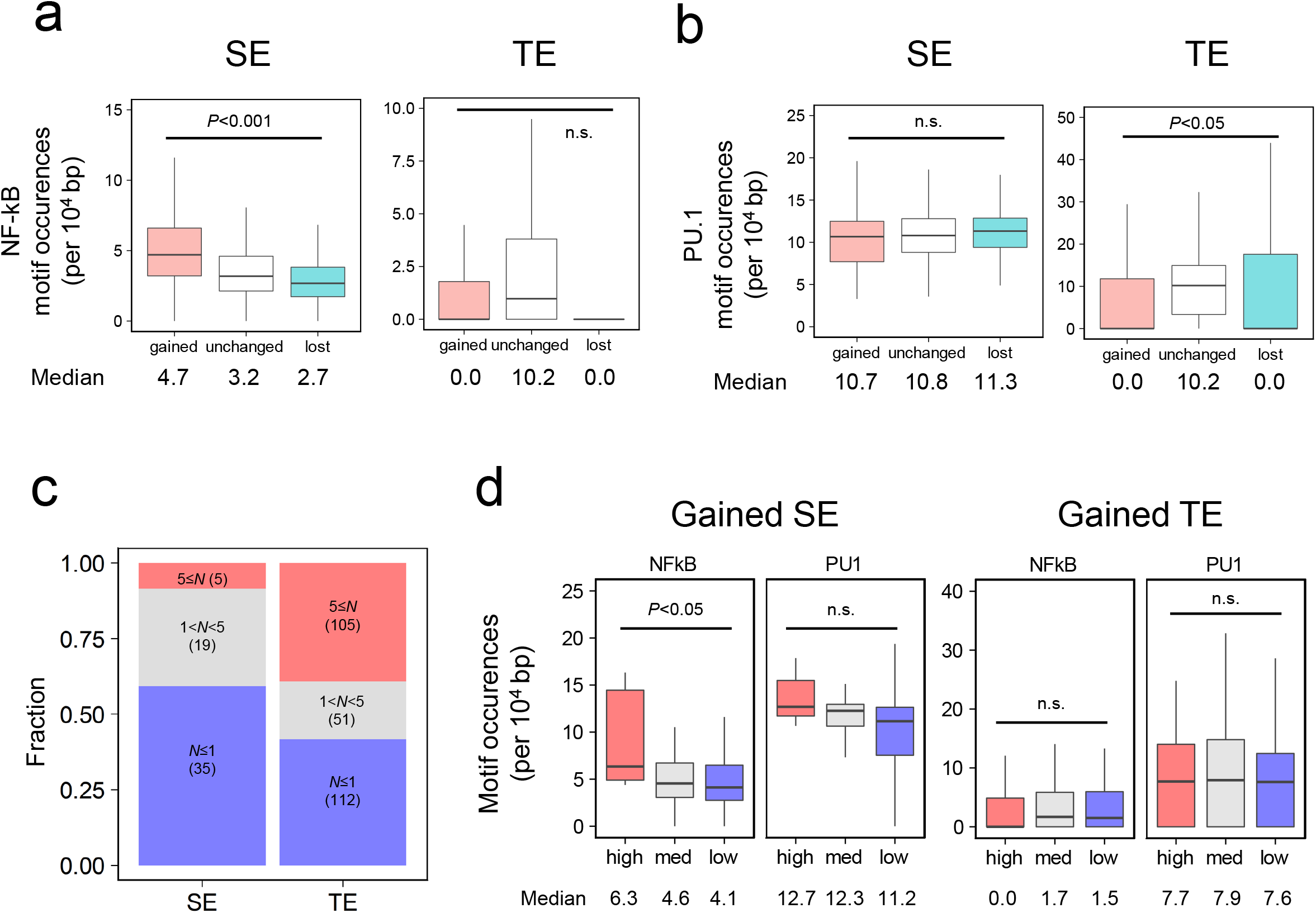
Motif and Hill function analysis. (a-b) Motif occurrences of (a) NF-κB and (b) PU.1 at SE and TE calculated using “findMotifsGenome.pl” program with “-find” option of Homer. Number of peaks: Gained SE, 280; Unchanged SE, 669; Lost SE, 280; Gained TE, 9024; Unchanged TE, 18050; Lost TE, 9021. P-value was calculated using one-way ANOVA with undersampling (n = 280), n.s.: not significant. (c) Bar plot of the Hill coefficient for mean gene expression across doses. Genes with Hill coefficient >9 and <0.3 were removed. (d) Boxplot of motif occurrences for NF-kB and PU.1 at gained SE and gained TE with categorized Hill coefficients. High, 5≤N<9; Med, 1<N<5; Low, 5≤N<9. *P*-values were calculated using one-way ANOVA with undersampling, except for gained SE in the high category (n = 19), n.s.: not significant.

SEs have exhibited switch-like gene expression defined by high dose-response Hill coefficient (*N*)^8,24^. We investigated this property by obtaining the Hill coefficient of gene expression across the anti-IgM concentration of our data. We performed optimization to the Hill function for the genes (0.2< fold change) (Supplementary Fig. 12), further categorized the genes based on the Hill coefficient (see Methods) (Fig. 4c). We further analyzed the motifs at the enhancers of these genes categorized by the Hill coefficient (Fig. 4d). The threshold gene expression in B cell has been reported to be tied to the number of NF-κB binding through RelA ChIP-seq and motif analysis^8^. Interestingly, we observed that motif density of NF-κB significantly increased across Hill coefficient in gained SE (6.3 for high, 4.6 for mid, 4.1 for low), albeit not TE. In contrast, PU.1 motif density at enhancers did not present significant changes across different Hill coefficients (12.7 for high, 12.3 for mid, and 11.2 for low). *CD83* and *NFKBIA* contained 5.9 and 10.6 NF-κB motifs per 10^4^ bp in their SE regions (Supplementary Fig. 13). The result is consistent with that *NFKBIA* exhibited bimodal distribution while that of *CD83* is monotonic (Fig. 2f-g, Supplementary Fig. 7). While the fitting of *N* based on mean expression is rough due to some genes having bimodal distribution (especially *NFKBIA)* (Supplementary Fig. 12), our result confirmed that the number of NF-κB binding to SE defines the properties of switch-like expression. The result also suggests that the local concentration of NF-κB in the nucleus strongly affects gene expression patterns.

### Gene expression heterogeneity is induced by gained cis-regulatory interactions

Further, we focused on the TEs and SEs in DEGs between activated and inactivated cells to examine the transcriptional heterogeneity. We observed that SE-associated genes have a higher Fano factor at all anti-IgM concentration ranges (Fig. 5a) and that this trend is proportional to the stimulation dose (Fig. 5b), as compared with TE. Further, there is a significant change in the Fano factor increased across different enhancer categorizations, where gained SE has a significantly higher Fano factor (Supplementary Fig. 14). Fano factor change between 10 μg/mL and 0 μg/mL is correlated with the increase in ATAC-signal (*R*=0.49 for SE, *R*=0.19 for TE) (Fig. 5c), thus, an increase in chromatin accessibility at enhancers are followed by an increase in heterogeneity in gene expression. In contrast, the classification of genes into high and low Fano factors does not present agreement with Hill coefficient classification (Supplementary Fig. 15).

**Figure 5.**
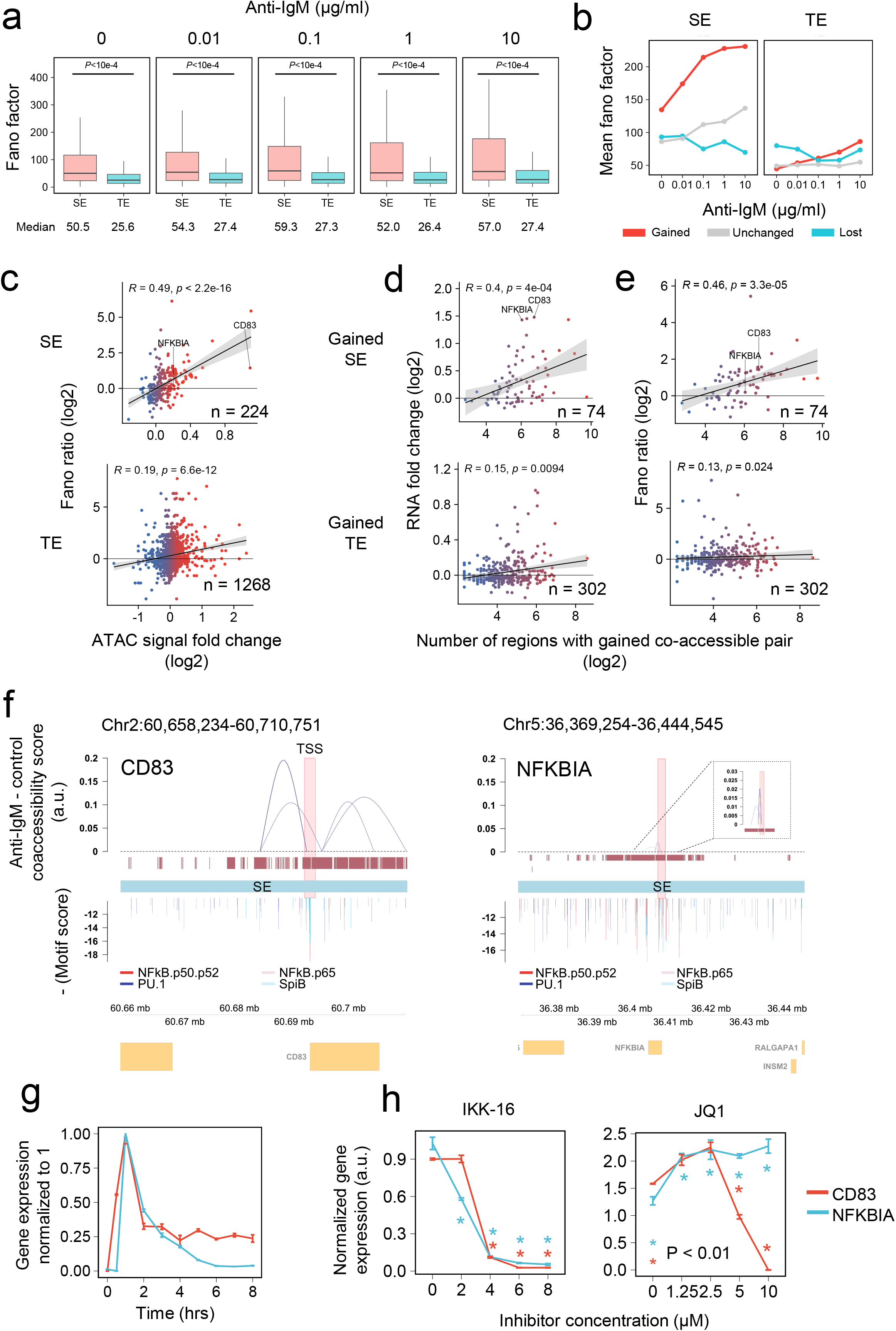
SE relationship with heterogeneity in gene expression. (a) Boxplot of Fano factor at various doses for DEGs associated with gained, lost, unchanged TE and SE. Number of genes: SE, 224; TE, 647. *P*-value was calculated using Welch’s unpaired t-test with undersampling (n = 224). (b) Line plot of mean Fano factor change across dose for SE and TE. Number of genes: Gained SE, 82; Unchanged SE, 100; Lost SE, 42; Gained TE, 242; Unchanged TE, 384; Lost TE, 21. (c) Correlation plot of Fano factor ratios (10 vs 0 μg/mL anti-IgM-stimulated cells) against ATAC intensity fold changes at SEs and TEs of marker genes. *R* = spearman correlation. (d) Correlation plot of RNA fold changes (activated vs inactivated cells) against numbers of co-accessible pairs. R = spearman correlation. (e) Correlation plot of Fano factor ratios (10 vs 0 μg/mL anti-IgM-stimulated cells) against numbers of co-accessible pairs. *R* = spearman correlation. (f) Co-accessibility score differences obtained using Cicero between stimulated and unstimulated cells shown between genomic regions interacting with regions ±1 kb around the annotated start site of *CD83* (left) *and NFKBIA* (right). (g) Time-course RT-qPCR of *CD83* and *NFKBIA* upon stimulation with 1 μg/mL anti-IgM. Gene expression was normalized to *GAPDH*. Error bar = SD. (h) RT-qPCR results of IKK-16 (left) and JQ1 (right) treatment 60 min prior to stimulation with 1 μg/mL anti-IgM for 60 min (N = 3). Gene expression was normalized to *GAPDH*, and *P*-values were calculated using Student’s unpaired t-test against dose 0 for each dose point. Error bar = SD.

Due to the complex arrangement of high-order chromatin structures involving SE-mediated gene expression, it remains difficult to generalize the contribution of chromatin accessibility of SEs to transcriptional regulation^40^. Therefore, we utilized single-cell ATAC-seq data for co-accessibility analysis using Cicero^25^ to elucidate the putative chromatin contact between enhancers and genes. We predicted accessible genomic regions that are potentially in close physical proximity by determining their co-accessibility scores and changes upon 10 μg/mL anti-IgM stimulation. Co-accessibility calculations were performed separately for the stimulated and unstimulated samples. The peaks used for co-accessibility analysis were obtained by merging shortly stitched peaks (108 bp) from both samples. Regions were filtered based on 0.1≤ co-accessibility scores in the stimulated samples and 0.05≤ in differences between the stimulated and unstimulated samples to detect significant increases in genomic interactions. To determine whether co-accessibility is correlated with gene expression amplitude and heterogeneity, we annotated each gene with co-accessible pairs above the threshold and observed a positive correlation between the number of regions with co-accessible pairs and their RNA fold changes (Fig. 5d) or Fano factor ratios (Fig. 5e). RNA fold-changes (*R*=0.4 for gained SEs, *R*=0.15 for gained TEs) and Fano factor (*R*=0.46 for gained SE, *R*=0.13 for gained TE) were positively correlated with a number of the regions with gained co-accessible pairs for SEs. These results suggest that the increased number of cis-regulatory genomic interaction is a common mechanism for enhanced gene expression and transcriptional heterogeneity.

To closely examine *CD83* and *NFKBIA*, we visualized the co-accessible region pairs within thresholds between peaks residing ±1 kb of each gene annotated transcriptional start site (TSS) and other positions in the same chromosome (Fig. 5f, Supplementary Fig. 16). Surprisingly, no co-accessible regions within annotated SE paired with the peaks ±1 kb of the *NFKBIA*-annotated TSS were observed above the 0.05 score threshold, while several regions associated with *CD83* were observed (Fig. 4d–e). Stronger and more long-range interactions were gained for *CD83* compared to *NFKBIA* (Supplementary Fig. 16). These results indicate the increase in the formation of higher chromatin interactions upon cell activation for *CD83* than *NFKBIA*, and this contributes to more enhanced expression heterogeneity.

Finally, we performed time-course RT-qPCR analysis on these two genes in the presence of IKK inhibitor and JQ1 to investigate their dependence on the NF-KB pathway or requirement of BRD4 for their SE formation. *CD83* expression pattern revealed a sustained pattern, while *NFKBIA* expression revealed a transient pattern on both qPCR and microarray data^41^ (Fig. 5g, Supplementary Fig. 17), suggesting that the differences in transcriptional regulation between these genes are potentially biologically different (*NFKBIA* as a feedback regulator of signaling pathway vs. *CD83* as a B cell differentiation marker). Further, we confirmed that both genes were NF-κB pathway-regulated since the IKK-16 treatment suppressed the expression of both genes (Fig. 5h left). In contrast, JQ1 treatment significantly suppressed *CD83* expression proportionally with higher JQ1 concentration, while *NFKBIA* expression slightly increased (Fig. 5h right), suggesting that *CD83* gene expression was more highly controlled by BRD4, but *NFKBIA* was not. While we speculate that this difference is caused secondary to the initial accessibility in the enhancer region of *NFKBIA* in the absence of stimuli (Fig. 3a) and that the chromatin accessibility change was comparably small upon stimulation (Fig. 2e), there might be other unknown mechanisms causing the increase in gene expression which have also been reported previously^42^.

## Discussion

In this paper, we aimed to reveal the SE activation and transcriptional regulatory mechanism controlled by NF-κB transcription factor using live-cell imaging and single-cell gene expression and chromatin accessibility analyses. We used DT40 B cells, which were derived from chicken bursal lymphoma. Previous reports have revealed that the BCR-NF-κB activation pathway and its dynamics in mouse primary B cells are reproducible in DT40 cells^7,43^. We have also demonstrated a consensus in SE classification for the target genes such as *CD83* and *NFKBIA*, which are expressed at slightly different timing after anti-IgM stimulation (Supplementary Fig.17), in these analyses regardless of the difference in analytical approaches (H3K27ac ChIP-seq vs. ATAC-seq) (Supplementary Fig. 9a).

Initially, we characterized RelA foci properties through fluorescence imaging analysis and observed some properties, such as sensitivity to LLPS perturbation, BRD4 dependence, and positive cooperativity. These analyses thus revealed that NF-κB nuclear aggregates exhibited properties similar to those of well-known SE condensates^19,20,24,44^.

SE-mediated gene regulation has been reported to cause transcriptional heterogeneity or noise^23^. However, another report presented that transcriptional noise is reduced in SE-mediated gene regulation^24^. Through experimental and computational approaches, we revealed that NF-κB-mediated SE formation strongly modulated transcriptional heterogeneity. We observed that genes with an increased heterogeneity upon increasing stimulation dose are enriched with cell-type-specific immune regulatory genes (Supplementary Fig. 5b), supporting a previous report where heterogeneity in gene expression is tied to biological functions and may be used by cells as a bet-hedging or a response distribution mechanism^45,46^, where cells exhibit heterogeneity to enable response to changing environment and also allowing dose-dependent fractional activation respectively. This was observed in *CD83*, a B cell activation marker, demonstrating the involvement of heterogeneity in B cell development.

As a mechanism to produce gene expression heterogeneity in phenotypically identical cells, we observed that co-accessibility, which has been observed to be concordant with genomic contacts, is correlated to Fano factor change, indicating that gene expression heterogeneity possibly stems from cis-regulatory genome interactions. NF-κB activation has been reported to be tied to the increase in heterogeneity in some genes and is attributed to the accumulation of Ser5p RNA Polymerase II^21^, which has also been reported to accumulate at enhancer regions^47^. This accumulation of RNA Polymerase II is proposed to assist in gene expression activation through enhancer-promoter contact^48^. Our results support these conclusions since co-accessibility or putative cis-regulatory interactions correlate to Fano factor changes. However, we also observed a weak correlation, which may be caused by the inequality of different enhancer elements, such as presenting additive properties reported in gene activation in *Drosophila* development^49^. The enrichment of TATA motif has also been proposed to generate transcriptional heterogeneity^23^. However, we observed a higher occurrence of TATA box in genes associated with lost SE (Supplementary Fig. 18) which might have caused gene expression heterogeneity in unstimulated cells. This heterogeneity might be owing to the difference in Pol II loading intervals^50^, however the noise associated with gained SE is possibly generated by the fluctuation of high-order biomolecular assembly and thus the source of heterogeneity in gene expression might be different. While SE can form phase-separated transcription hubs containing multiple enhancers and/or promoters that might enable the higher diffusion rate of active enhancers and consequently, higher possibility of genomic DNA interactions^51^, the current analytical method to analyze SE ignores individual enhancer properties^52^ and non-linear interactions. Therefore, a dissection of each enhancer contact may be required to understand how these differences arise in the context of SE.

To answer this question, we tried to utilize a CRISPR-Cas9 system with double-flanking gRNA and enhanced specificity *S. pyogenes* Cas9^53^ to induce long deletion in regions with the highest co-accessibility score change in the *CD83* promoter (Fig. 5). Unfortunately, we were unable to confirm the deletion of the regions within *CD83* SE, which may be attributable to the possibly incomplete genome database of the chicken reference genome used for the gRNA design (GRCg6a). We were able to induce a long deletion in the enhancer region outside of the *CD83* SE. However, the obtained cell clone demonstrated low survival and impaired proliferation rates. Thus, we were unable to confirm the changes in gene expression dynamics upon deletion of this enhancer region.

In contrast, SE has been reported to induce both switch-like gene expression^8^ and enhanced heterogeneity^23^. Mathematical model has been used to explain that the threshold response is generated by the number of multivalent interactions mediated by transcriptional assembly within SE^24^. Therefore, genes showing threshold response might also show higher gene expression heterogeneity. However, our analysis revealed that this is not the case. Although we observed that there is a higher proportion of genes with Fano factor above the median in genes with higher Hill coefficient (1<*N*<5), there is no clear relationship between heterogeneity and threshold gene expression (Supplementary Fig. 15). These results indicate that although enhanced gene expression and transcriptional heterogeneity are controlled by genomic contacts between chromatin accessible regions, switch-like expression of genes is controlled by NF-κB binding to SE (Fig. 4) and those contributions are different for an individual gene. With this regard, we believe that the velocity and duration of nuclear translocation of NF-κB, along with interaction with other transcriptional factors might strongly affect those transcription responses.

Lastly, we noticed that sequence-based identification of SEs may not truly reflect the biologically equal functions of SEs for each target gene. For example, *NFKBIA* and *CD83* mRNAs exhibited varying sensitivities to JQ1 (Fig. 5h, right). Since the *NFKBIA* SE had constitutively open chromatin prior to anti-IgM stimulation (Fig. 3c), the signal-dependent effect on SE regulation of *NFKBIA* appeared to be small. These two genes demonstrated large fold-changes in gene expression upon SE regulation. However, the molecular level of SE regulation in each gene varied, which might have caused differences in transcriptional heterogeneity. Therefore, our results suggest that the functional evaluation of SE should not solely rely on sequence analysis, but also additional experimental methods.

In conclusion, these results provide an alternative insight into the mechanism of SE-mediated gene regulation upon NF-κB activation. Especially in the generation of biologically relevant gene expression, heterogeneity in phenotypically identical cell populations by SE is comprehended through gained cis-regulatory interactions. We expect similar methods to be applied in further studies on other signaling pathways regulated by SEs.

## Materials and methods

### DT40 cell culture

Wild-type and RelA-GFP-expressing DT40 chicken bursal lymphoma cells were obtained from Dr. Shinohara^7^. Mouse RelA-eGFP with eGFP on the C terminal was cloned into a pGAP vector containing Ecogpt resistance gene targeting endogenous GAPDH locus. This construct was further electroporated into wild-type cells and selected using Ecogpt to produce RelA-GFP-expressing DT40 cells^7^. The DT40 cells were cultured in RPMI-1640 without phenol red (Wako, Japan) supplemented with 10% fetal bovine serum (Sigma-Aldrich, USA), 1% (v/v) chicken serum (Nippon Bio-test Laboratories, Japan), 75 μM 2-mercaptoethanol (Gibco, USA), 1 mM sodium pyruvate (Wako, Japan), 1% (v/v) penicillin-streptomycin solution (Nacalai Tesque, Japan), 1% (v/v) 100x MEM non-essential amino acids solution (Wako), and 2 mM L-glutamine (Nacalai Tesque). The cells were cultured at 39°C and 5% CO_2_ in a humidified incubator.

### mKate2-BRD4S transposon plasmid construction

We engineered the PB-TA-ERP2-mKate2-BRD4S construct from two addgene clones (65378^54^ and 80477^55^) and pmKate2-H2B (Evrogen, Russia). Overlap extension PCR was performed to amplify the mKate2 insert from the mKate2-H2B plasmid while adding the attB1 adapter and linker sequence. Another round of overlap extension PCR was performed to amplify the BRD4S insert from GFP-BRD4 while adding the attB2 adapter and linker sequence. A final round of fusion PCR was performed to fuse the fragments containing mKate2 and BRD4 to create an insert containing mKate2-linker-BRD4S. The primers used for the overlap extension PCR are listed in Supplementary Table 3. A BP reaction using BP Clonase II (Invitrogen, USA) was performed to clone the insert into pDONR221 (Invitrogen), creating the entry clone pENTR221-mKate2-BRD4S. The entry clone was amplified using NEB-Stable (New England Biolabs, USA). An LR reaction using LR Clonase II (Invitrogen) was performed with the destination vector PB-TA-ERP2^55^ to create the final expression vector.

### Cell transfection

Cell transfection was performed using the Neon Transfection System (Invitrogen). A total of 2 μg of plasmid was mixed with 1 × 10^6^ cells in 10 μL buffer R. Electroporation was performed at 1400 V for 30 ms with 1 pulse. For piggyBac co-transfection, a mass ratio of 1:3 (PB:pBase) of the plasmid was used.

### Quantification of RelA foci in single cells

Prior to stimulation and subsequent live imaging, DT40 cells were concentrated 5× into a serum-free medium and incubated in a 39 °C incubator with 5% CO_2_. Cells were further transferred into an eight-welled coverglass chamber (Iwaki, Japan). An inverted microscope IX81 (Olympus, Japan) equipped with a CSU-X1 confocal scanner unit (Yokogawa, Japan) and oil-immersion objective (100×, NA 1.45) was used to obtain images. MetaMorph software (Molecular Devices, USA) was used to obtain 13 z-slices of stack images at 1 μm increments in the z-direction. The image resolution used was 512 × 512 pixels (1 pixel = 0.16 μm). The observation chamber was maintained at 39 °C during observation. FIJI ImageJ 1.52i (https://imagej.net/Fiji/Downloads) was used to count the foci using a custom macro, where the diameter for foci detection was set to 0.96 μm.

### JQ1, IKK-16, and 1,6-hexanediol cell treatment

For JQ1 and IKK-16 inhibition, the DT40 cells were suspended in 5 μM JQ1 (Selleckchem, USA) and 6 μM of IKK-16 (Selleckchem) 60 min before anti-IgM stimulation. For 1,6-hexanediol inhibition, 5% of 1,6-hexanediol (Sigma-Aldrich) was added after anti-IgM stimulation.

### Quantification using smRNA-FISH

Fluorescence-conjugated chicken *GAPDH, CD83*, and *NFKBIA* probes were generated for smRNA-FISH using the Stellaris Probe Designer (https://www.biosearchtech.com/support/tools/design-software/stellaris-probe-designer) (Biosearch Technologies, UK) according to the protocols of Biosearch Technologies. The sequences of the probes are presented in Supplementary Table 4. A total of 1 × 10^7^ DT40 cells in 600 μL RPMI without supplements were stimulated with anti-IgM or PBS (control) for 30 min before washing with PBS and resuspending in 1 mL suspension buffer. Procedures following cell fixation were performed as described in the manuals of Biosearch Technologies for suspension cells. The fixed cells were mounted using Vectashield (Vector Laboratories, USA), sandwiched in cover glasses (Matsunami Glass, Japan), and sealed with clear nail polish prior to imaging.

A DeltaVision Elite - Olympus IX71 (Olympus) fluorescence microscope equipped with a Coolsnap HQ2 camera (Photometrics, USA) and an oil-immersion objective (60×, NA 1.42) was used for image acquisition. SoftWoRx software (Applied Precision, USA) was used for image acquisition and deconvolution. FISH-quant v3 was used to quantify the RNA-FISH images^56^. The default parameters were used in the quantification.

### Quantitative RT-PCR (qRT-PCR) analysis

Total RNA was collected from the DT40 cells using a NucleoSpin RNA kit (Macherey-Nagel GmbH & Co., Germany) and subjected to complementary DNA synthesis and quantitative PCR using a ReverTra Ace qPCR RT Kit (Toyobo Life Science, Japan) and KOD SYBR qPCR kit (Toyobo Life Science) according to the manufacturer’s protocol. PCR cycling conditions were as follows: 40 cycles of 10 s at 98 °C, 10 s at 60 °C, and 30 s at 68 °C. The primers used for qRT-PCR are listed in Supplementary Table 5. Expression values (n = 3) were normalized to those of *GAPDH*. For the inhibitor treatment, IKK-16 and JQ1 were added 60 min before anti-IgM stimulation to investigate the dependences of *CD83* and *NFKBIA* on SEs.

### Single-cell RNA-sequencing analysis

The DT40 cells were stimulated with anti-IgM (0, 0.01, 0.1, 1, and 10 μg/mL) for 1 h and sorted using an SH800 cell sorter (Sony, Japan) with a 130 μm sorting chip to select single cells. RamDA-seq with oligo-dT primer single-cell RNA-seq method was used for cDNA preparation^31^. The samples were sequenced using HiSeq2500 (Illumina, USA).

A quality check of the data was performed using FastQC. Trimming was performed using TrimGalore (with Cutadapt v2.3; https://www.bioinformatics.babraham.ac.uk/projects/trim_galore/) with default options, and alignment to the *Gallus gallus* reference genome (GRCg6a) was done using STAR v2.7.1a with default options^57^. Further, a gene count table was obtained from the alignment files using featureCounts v1.6.4 with “-t exon -g gene_id” options and the annotation GTF file GRCg6a.96 from ENSEMBL (ftp://ftp.ensembl.org/pub/release-96/gtf/gallus_gallus/)^58^.

Seurat v3.2.1 was used for clustering and differential gene expression analysis of the scRNA-seq data^32^. Prior to clustering, a quality check of the data was performed to remove cell outliers (total count ≥1.5 million, detected genes ≥8500, and mitochondrial gene count ratio <0.04) resulting in 89, 92, 87, 92, and 93 cells used in the analysis of the 0, 0.01, 0.1, 1, and 10 μg/mL anti-IgM concentrations, respectively. Data normalization was performed using Baynorm^59^ and the log scaling method in Seurat. Data were regressed based on the cell cycle scoring with the “CellCycleScoring()” function of Seurat and mitochondrial gene count ratio. The top 2000 variable features were used for dimensionality reduction and clustering with a resolution of 0.2. DEGs were extracted using the “FindAllMarkers()” function of Seurat. Pseudo-time analysis was performed by creating a principal curve using the Princurve v2.1.4 R package on the dimensionality-reduced projection with the Lowess smoother.

### Single-cell ATAC-sequencing analysis

All protocols for generating scATAC-seq data on the 10x Chromium platform (10x Genomics, USA), including sample preparation, library preparation, and instrument and sequencing settings, are described below available here: https://support.10xgenomics.com/single-cell-atac. Prior to nuclei extraction, the DT40 cells were stimulated with 10 μg/mL of anti-IgM or PBS for 60 min. Cellranger-atac count v1.1.0 (https://support.10xgenomics.com/single-cell-atac/software/pipelines/latest/algorithms/overview) was used with default options for performing quality checks and mapping the scATAC-seq data to the genome. The sequence files were downsampled to 250 million reads before running the Cellranger-atac count pipeline. The reference genome used was the ENSEMBL genome with the annotation file GRCg6a.96 (ftp://ftp.ensembl.org/pub/release-96/gtf/gallus_gallus/).

### Identification of SEs and TEs

Peak calling and enhancer identification from ATAC-seq data were performed using Homer v4.10.4 (http://homer.ucsd.edu/homer/) using the bam files generated from the Cell Ranger pipeline. Tag directories were created for the bam file from each condition using the “makeTagDirectory” program with the “--sspe -single -tbp 1” option. Peak calling was performed using the “findPeaks” program with the “-style super -typical - minDist 5000 -L 0 -fdr 0.0001” option. This procedure stitches peaks within 5 kb and ranks regions by their total normalized number reads and classifies TE and SE by a slope threshold of 1. Peak annotation was subsequently performed using the “annotatePeaks.pl” program with the GRCg6a.96 annotation file. The consequent peak files were merged between each stimulation condition for the SE and TE peaks separately using the “mergeBed” program of bedtools. Peak annotation was performed for the second time for the merged peaks to create the final SE and TE peaks. ATAC fold-change was then calculated between both conditions for the merged peaks separately for SE and TE. Genes associated with both SE and TE were assigned only to the SE.

### ATAC-seq data visualization

Mouse primary splenic B-cell ATAC-seq data was obtained from the accession number DRA009931 of DDBJ ^8^. Mapping of data to produce BigWig files was performed using nf-core/rnaseq pipeline^60^ v3.3 (default options) and the default GRCm38 genome. Visualization of both primary and DT40 data was performed using Gviz v3.1.3 ^61^.

### Gene ontology analysis

Gene ontology analysis was performed using the function “enrichGO” of clusterProfiler v3.14.3 for gained SE and RNA upregulated genes^62^. The Ensembl gene id was converted to the mouse homolog gene id prior to enrichment analysis using the query “mmusculus_homolog_ensembl_gene”.

### Motif analysis

Analysis of motif occurrences was performed using Homer v4.10.4^63^ “findMotifsGenome.pl” program with “-find” option. “NFkB-p65-Rel(RHD)/ThioMac-LPS-Expression(GSE23622)/Homer,” “NFkB-p50,p52(RHD)/Monocyte-p50-ChIP-Chip(Schreiber_et_al.)/Homer” and “NFkB-p65(RHD)/GM12787-p65-ChIP-Seq(GSE19485)/Homer” were considered as NF-κB motifs. “PU.1-IRF(ETS:IRF)/BcellPU.1-ChIP-Seq(GSE21512)/Homer,” “PU.1:IRF8(ETS:IRF)/pDC-Irf8-ChIP-Seq(GSE66899)/Homer” and “PU.1 (ETS)/ThioMacPU. 1 -ChIP-Seq(GSE21512)/Homer” were considered as PU.1 motifs.

Analysis of motif occurrences at promoter regions was performed using Homer v4.10.4^63^ “findMotifsGenome.pl” program with “-find” option for regions spanning -300 bp and +50 bp from the annotated TSS. “TATA-Box(TBP)/Promoter/Homer” was considered as TATA-box motif.

Motif score calculation for enhancer regions was performed using FIMO v5.0.5 with the database obtained from Homer v4.10.4^63^. The motif files from Homer were converted using the “read_homer()” and “write_meme()” functions of universalmotif v1.4.10 R package.

### Co-accessibility analysis

Peak calling was performed for co-accessibility analysis in Homer v4.10.4 using the “findPeaks” program with the “-style super -typical -minDist 0 -L 0 -fdr 0.0001” option to identify peaks constituting SE and TE. The “FeatureMatrix()” function of Signac v1.0.0 was used to assign fragments from the “fragments.tsv” file previously filtered for cell barcoding to the bed file containing peaks.

Cicero v1.3.4.10^25^ was used to calculate the co-accessibility scores^25^ between the ATAC peaks using the reference genome GRCg6a.96. The “max_sample_windows” argument of the “distance_parameters()” function was set to 1000, and the “max_elements” argument of the “generate_cicero_models()” function was set to 500. The other options were set to default. Co-accessibility calculations were performed separately for both the stimulated and unstimulated cells. The final co-accessibility scores were determined as the differences between the co-accessibility scores of the stimulated cells and unstimulated cells, where the initial score was ≥0.1 in the stimulated cells. Gviz v1.30.3 was used to visualize the co-accessibility between genomic regions.

### Calculation of Hill coefficient

The focus dose-response curve was fitted to the Hill function with a constant additive term accounting for basal activity. The parameters were optimized using all methods of the “optimx()” function in the “optimx” R package, and the method with the best fit was selected^64^. The equation used was as follows:

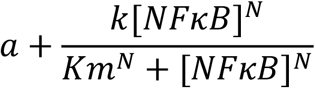

where *N*, *Km, k*, and *a* represent the Hill coefficient (cooperativity), the binding affinity of NF-κB to the enhancer region, the rate constant for foci formation, and the constant additive term accounting for basal activity, respectively.

For fitting of mean gene expression fold change to Hill coefficient in the Fig 4c-d, we performed optimization to the Hill function with removed basal activity term for genes with a fold change above 0.2 between 0 and 10 μg/mL anti-IgM and also removed genes with Hill coefficient above 9 and below 0.3 as these genes are either overfitted or do not fit at all. We then categorized genes with having high (5 < *N*), medium (1 < *N* < 5), and low (1 < *N*) Hill coefficients.

### Clustering of Fano factor change

Hierarchical clustering of Fano factor changes across anti-IgM doses was performed using the “hclust()” function of R with the method option “ward.D2” and the tree was cut at 4 branches.

### Statistics and reproducibility

The qPCR data are presented as the mean ± standard deviation (SD), wherein N = number of biological replicates. Data were evaluated using the unpaired Student’s t-test. The means were considered statistically significant at *p* < 0.05. Box plots represent the median (centerline), interquartile range (IQR; box limits), and 1.5× IQR for the whiskers.

## Data availability

All sequencing data were deposited in the DNA Data Bank of Japan (DDBJ) under the accession number DRA012330. The codes used for the bioinformatics analysis and imaging analysis are available at https://github.com/okadalabipr/Wibisana2021. Other data are available from the corresponding author upon request.

## Acknowledgements

We thank Dr. Kazuo Yamashita and Dr. Masakazu Ishikawa of KOTAI Biotechnologies, Inc. (Suita, Osaka, Japan) for preparing the scATAC-seq samples. We thank Dr. Akira Imamoto and Dr. Shinpei Kawaoka for their discussions regarding plasmid construction and CRISPR-Cas9 experiments as well as Dr. Takeya Kasukawa and Dr. Keita Iida for discussions regarding NGS analysis. M.O. was supported by JSPS KAKENHI grants 17H06299, 17H06302, and 18H04031 as well as JST-Mirai Program grant JPMJMI19G7. Y.S was supported by JSPS KAKENHI grant 19K22404. I. N. was partially supported by JST CREST grant JPMJCR16G3 and JPMJCR1926. This research was partially supported by the Platform Project for Supporting in Drug Discovery and Life Science Research (Platform for Drug Discovery, Informatics, and Structural Life Science) from the Japan Agency for Medical Research and Development (AMED). J. N. W. was supported by the Honjo International Scholarship Foundation.

## Author contributions

J.N.W. performed computational data analysis of scRNA-seq and scATAC-seq, conducted smFISH experiments and constructed the BRD4S-mKate2 cells. J.N.W., T.I., and Y.S. performed the imaging analyses. H.S. and N.Y. constructed the RelA-GFP-expressing cells and prepared the samples for RamDA-seq. T.H., M.U., M.E., and I.N. performed the single-cell sorting and RamDA-seq. J.N.W., T.I., Y.S., and M.O. interpreted the analysis results. J.N.W. and M.O. wrote the manuscript. Y.S. and M.O. conceived the study and performed overall direction and planning.

## Competing interests

The authors declare no competing interests.

## Notes

### Competing Interest Statement

The authors have declared no competing interest.

