## Supplementary figures and tables for "Enhanced transcriptional heterogeneity mediated by NF-κB super-enhancers"

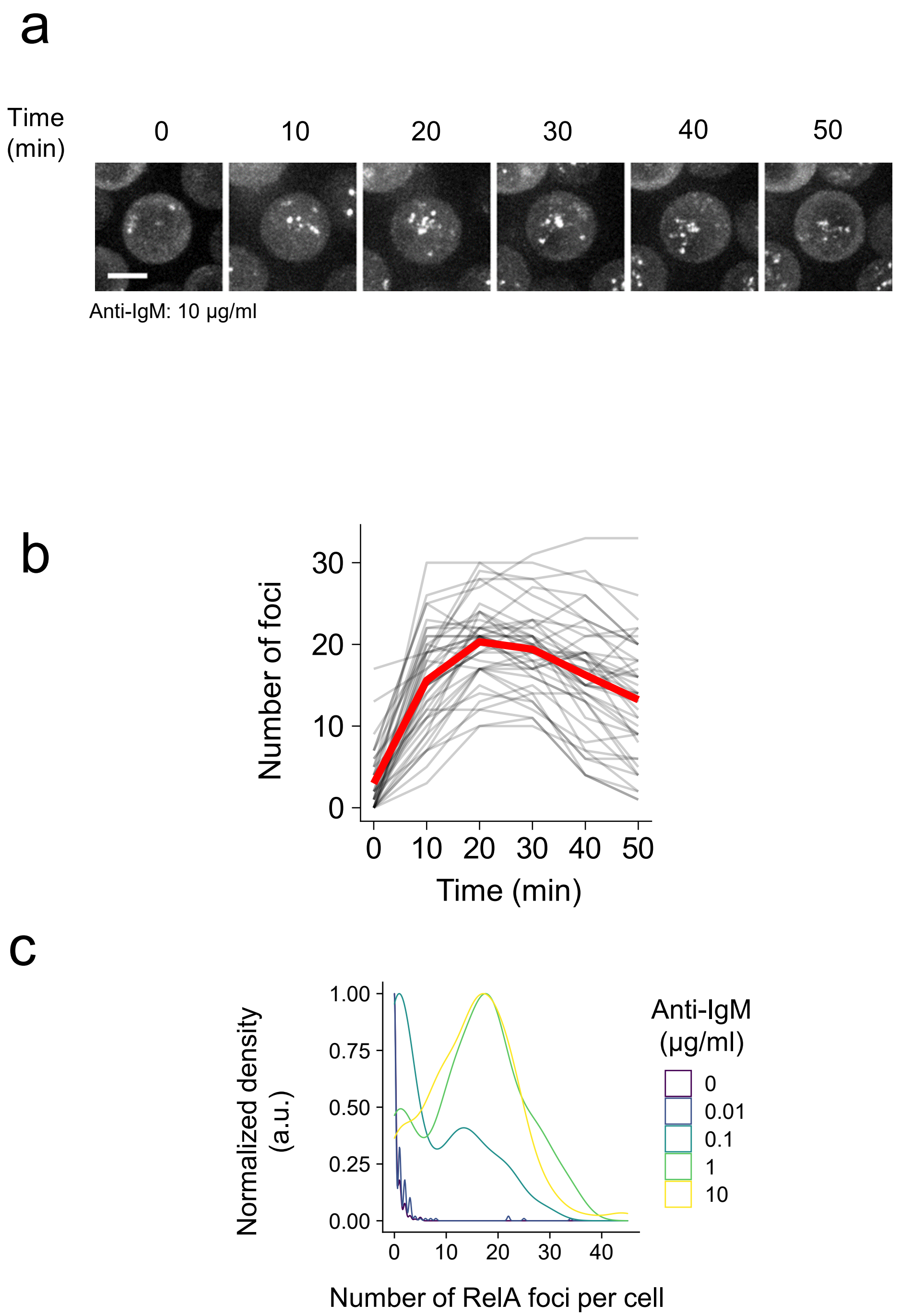


**Supplementary Figure 1. Dynamics of RelA nuclear translocation upon anti-IgM stimulation on DT40 cells.** (a) Representative fluorescence micrographs of a single cell upon stimulation with 10 µg/ml anti-IgM (scale bar, 10 µm). (b) Changes in the number of foci detected in single-cells across time upon addition of 10 µg/ml anti-IgM (n = 49). Red line indicates mean. (c) Distribution of RelA foci per cell upon stimulation with various doses of anti-IgM for 20 min.

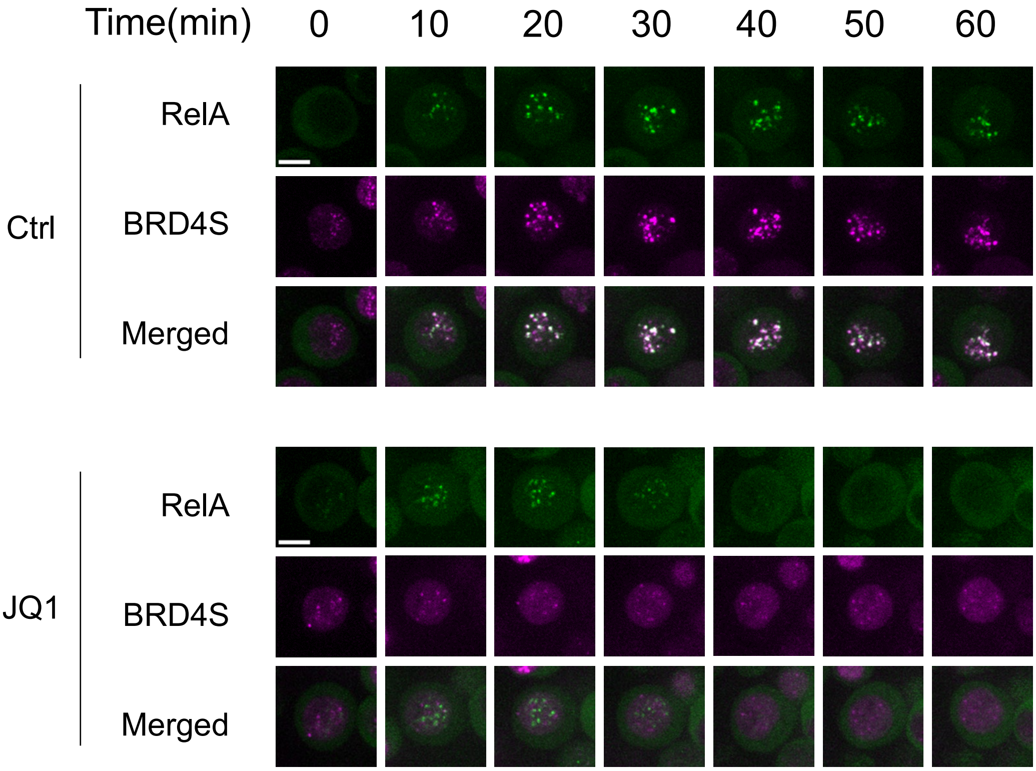


**Supplementary Figure 2. JQ1 treatment of mKate2-BRD4S and RelA-GFP coexpressing cells.** Time-lapse fluorescence micrographs of DT40 cells co-expressing mKate2-BRD4S and RelA-GFP upon stimulation with 10 µg/ml anti-IgM and pre-treatment with JQ1 (5 µM) for 60 min (scale bar, 5 µm).

a

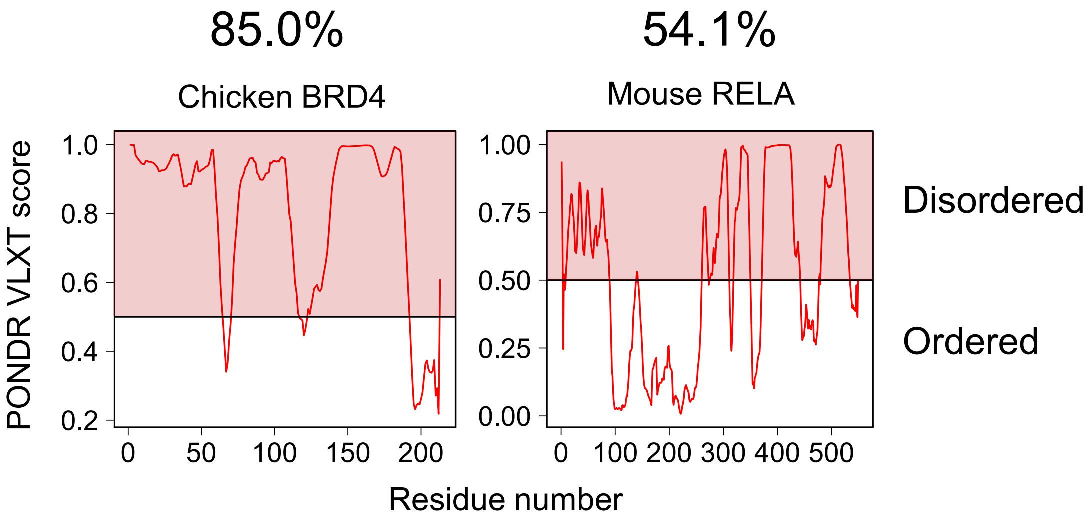


b

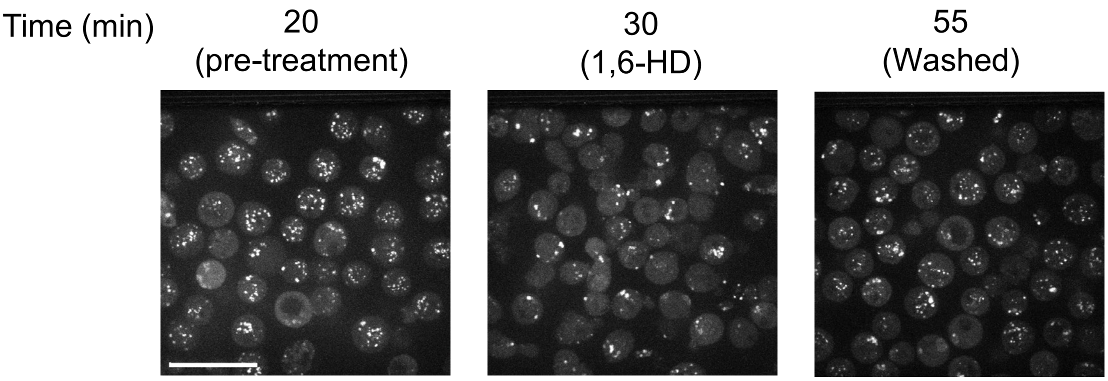


**Supplementary Figure 3. RelA-GFP foci demonstrate LLPS condensate-like biophysical properties.** (a) PONDR VLXT disorder scores of BRD4 and RelA. PONDR score more than 0.5: BRD4, 85.0%; RelA, 54.1%. (b) Representative fluorescence micrographs of a cell population stimulated with 10 μg/ml anti-IgM before treatment, after 1,6-hexanediol treatment, and upon washing (scale bar, 25 μm).

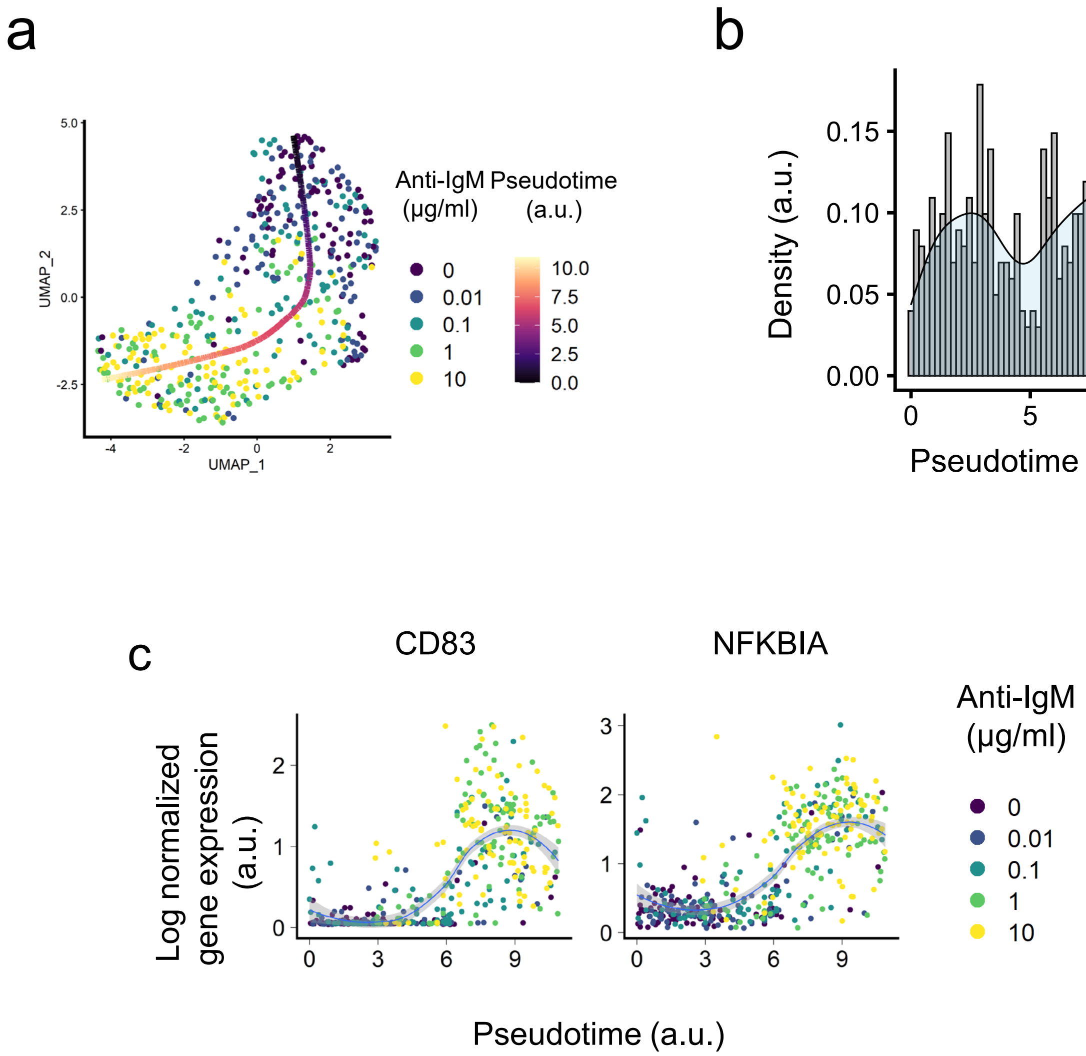


**Supplementary Figure 4. scRNA-seq analysis of DT40 cell stimulated with various doses of anti-IgM.** (a) Pseudo-time axis taken using a principal curve over a UMAP projection showing the doses of anti-IgM. (b) Distribution of cells across pseudo-time. (c) Gene expression of CD83 and NFKBIA in single-cells across pseudotime.

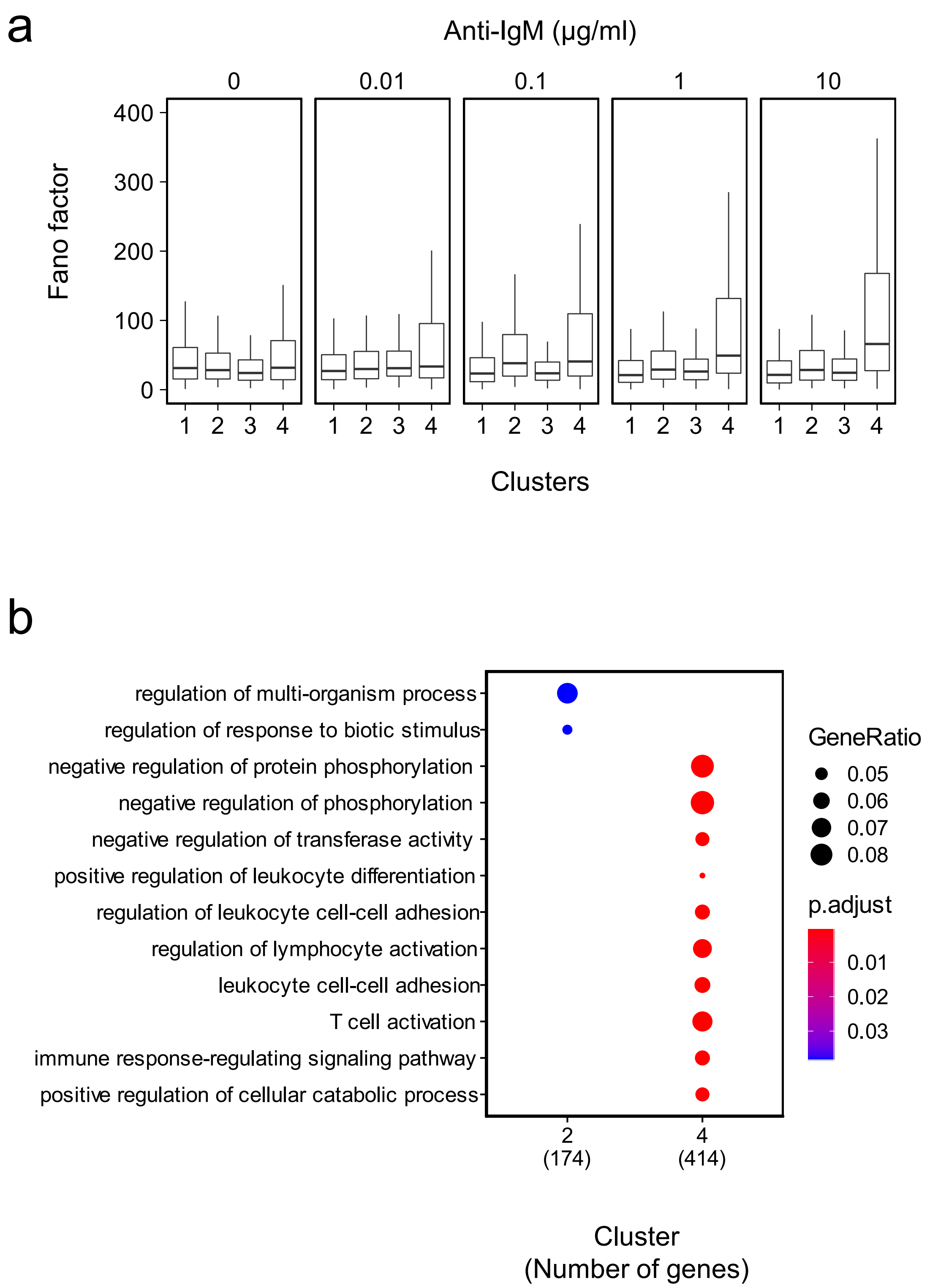


**Supplementary Figure 5. Analyses of genes in different heterogeneity clusters.** (a) Boxplot showing Fano factor of each clusters across different anti-IgM concentrations. (b) Biological processes gene ontology enrichment analysis for DEGs clustered according to Fano factor changes across anti-IgM doses.

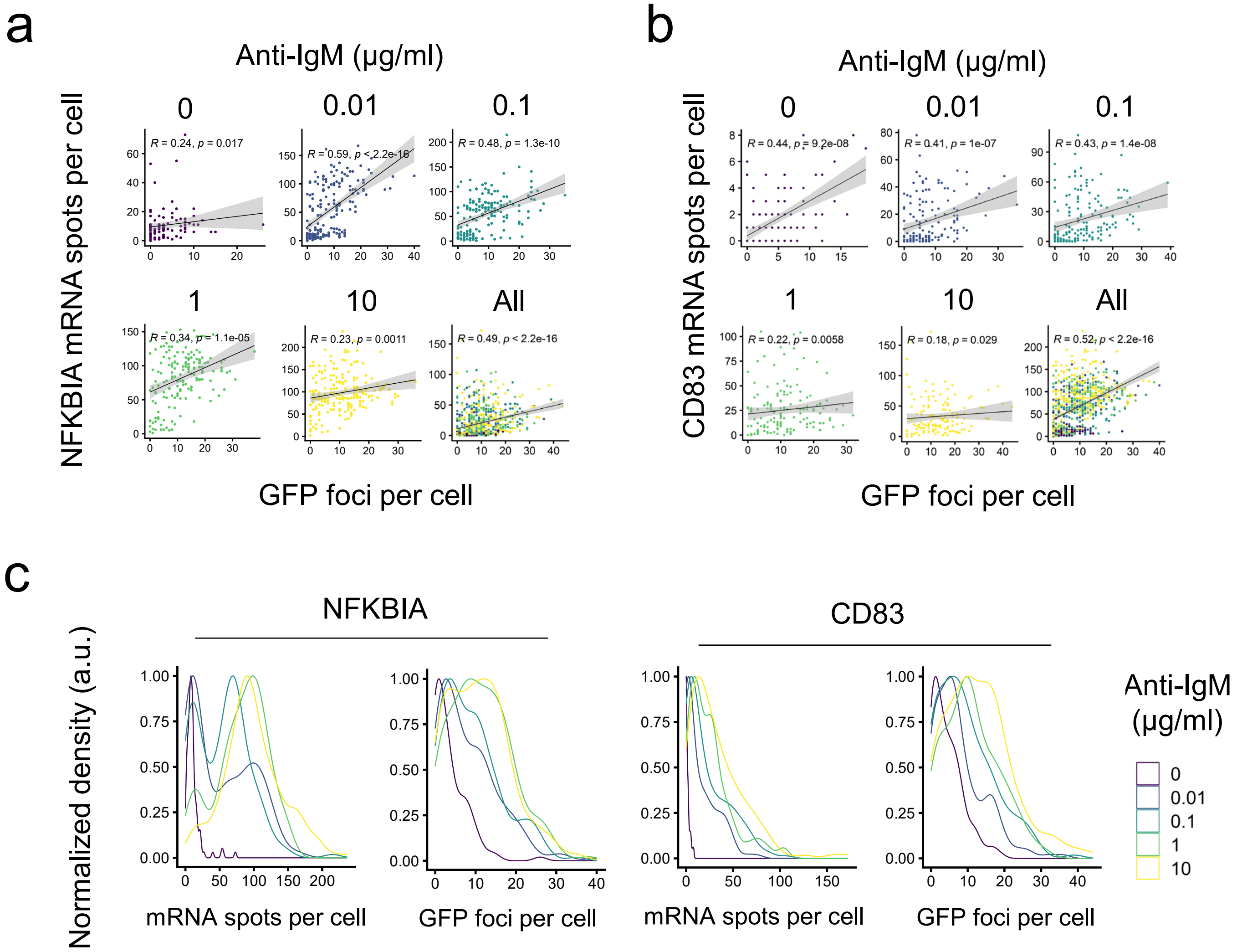


**Supplementary Figure 6. smRNA-FISH analysis of DT40 cells stimulated with various doses of anti-IgM.** (a–b) Correlation plot between GFP foci and mRNA spots per cell for various doses of anti-IgM. (c) Density plot of RNA spots and GFP foci at various anti-IgM doses from smRNA-FISH analysis.

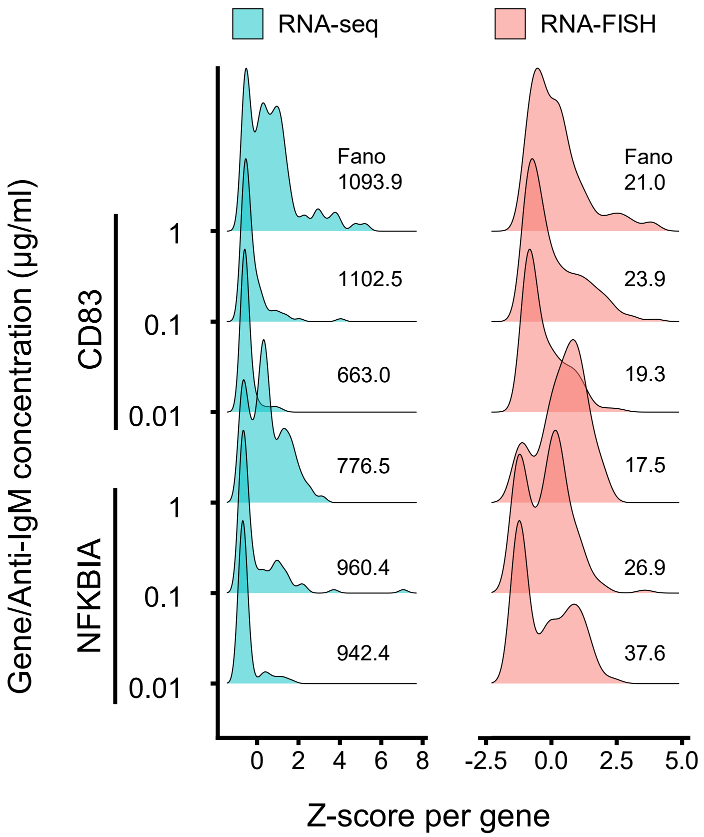


**Supplementary Figure 7. Gene expression distribution of *CD83* and *NFKBIA*.** Single-cell expression of *CD83* (B cell activation marker) and *NFKBIA* (NF-κB target gene) obtained from scRNA-seq and smRNA-FISH at various doses of anti-IgM.

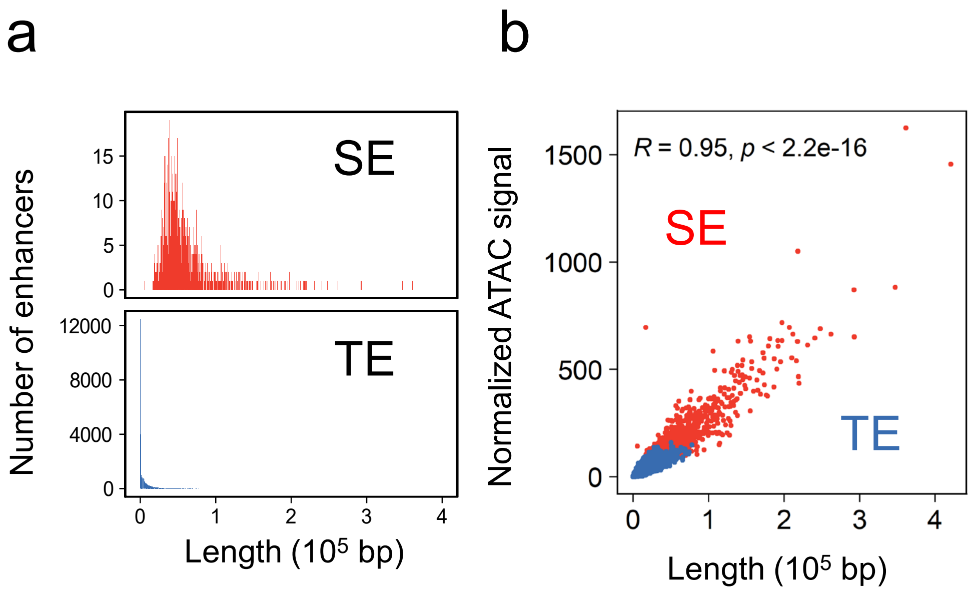


**Supplementary Figure 8. SE regions show longer length which correlates with ATAC signal.** (a) Length of classified SE and TE. (b) Correlation plot between enhancer length and normalized ATAC signal. R = pearson correlation coefficient.

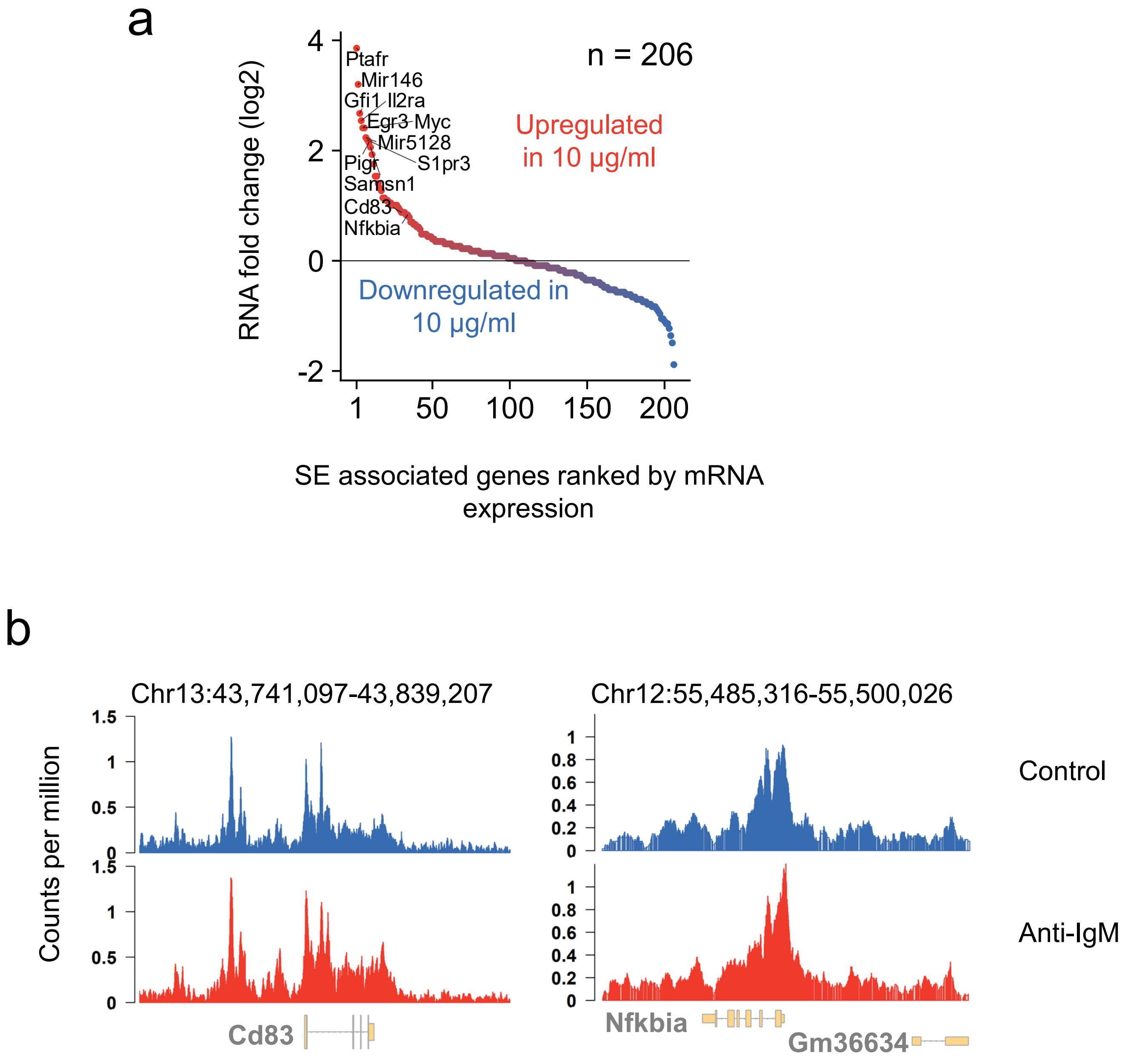


**Supplementary Figure 9. SE in primary B cells.** (a) Scatter plot of the mean fold-change of SE-associated genes between 10 µg/ml anti-IgM stimulated and control cells (Michida et al., 2020). Note that SE was identified from H3K27Ac ChIP-seq data (b) Track view of Cd83 and Nfkbia ATAC-seq data (Michida et al., 2020).

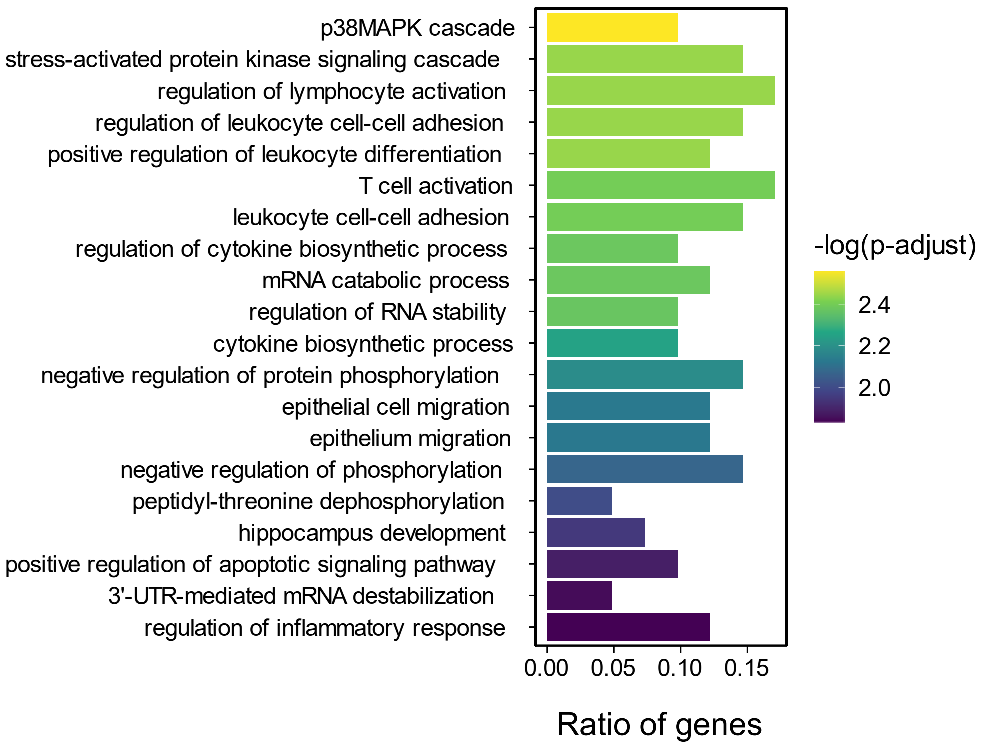


**Supplementary Figure 10. Gained SE upregulated genes show immune related biological functions.** Biological processes gene ontology (GO) enrichment analysis of 52 genes with both gained SE (upper quantile) and upregulated RNA (upper quantile).

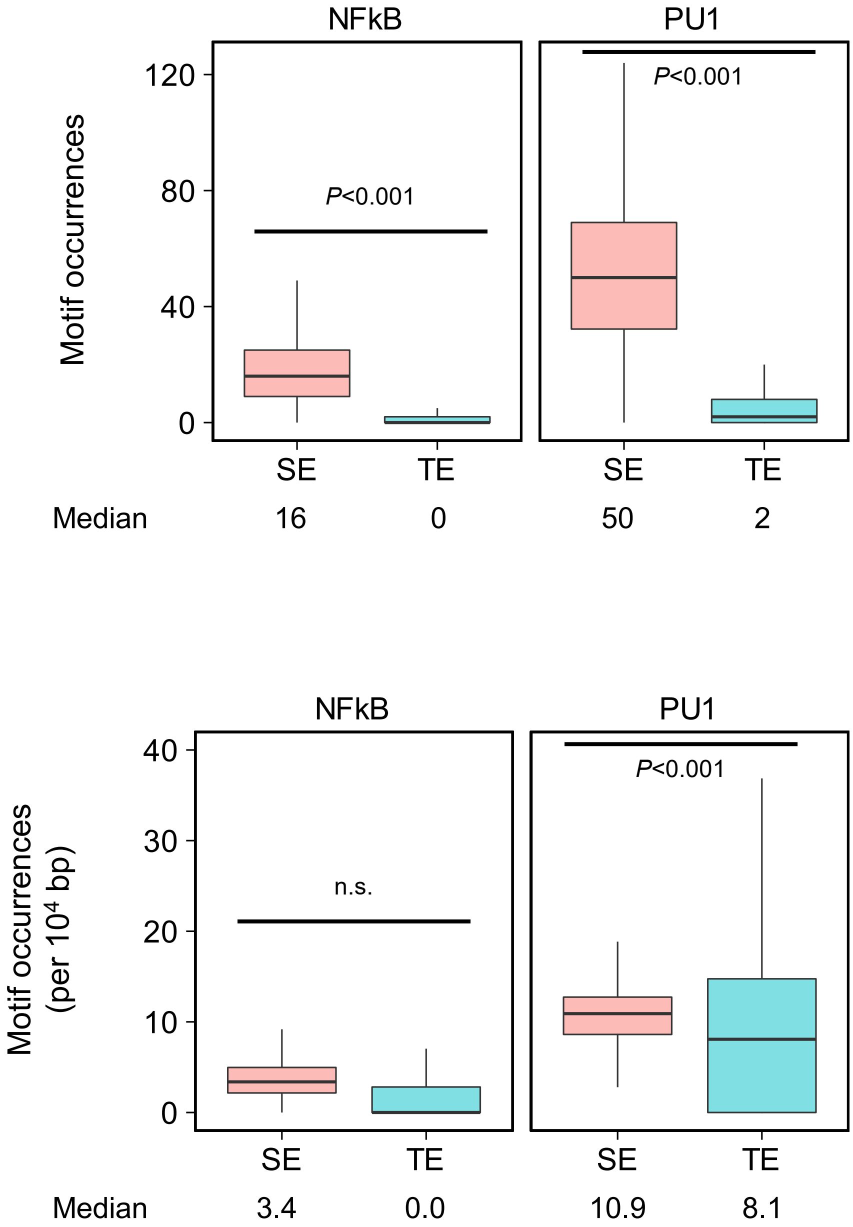


**Supplementary Figure 11. Motif occurrences of NF-kB and PU.1 at enhancer regions are higher at SE than TE.** Motif occurrences of both NF-kB and PU.1 at TE (37,686 peaks) and SE (1,118 peaks) calculated using "findMotifsGenome.pl" program with "-find" option of Homer. The *P*-values were calculated using Welch's t-test after undersampling (n = 280), n.s.: not significant.

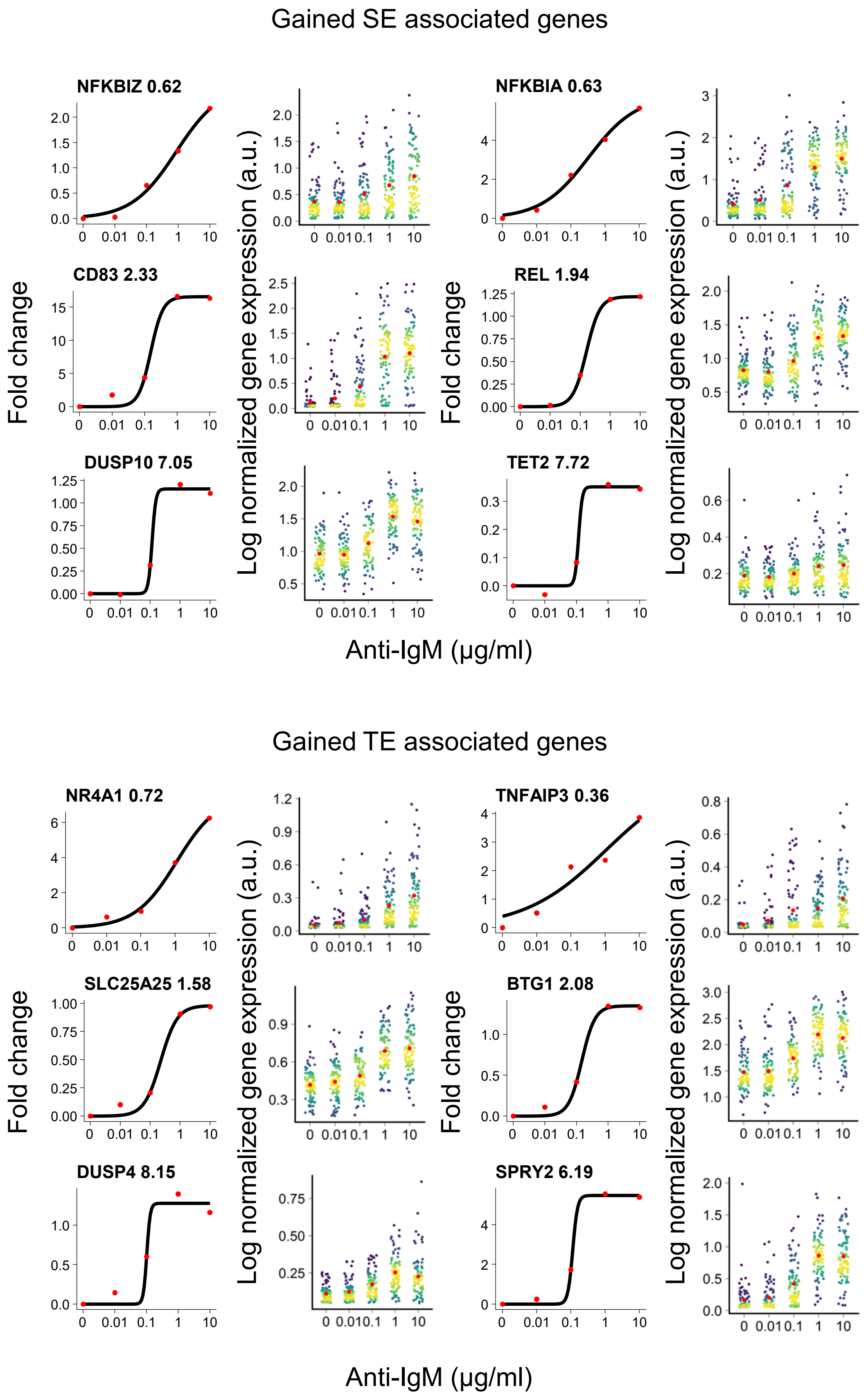


**Supplementary Figure 12. Fitting of mean gene expression to Hill function for gained SE and TE associated genes.** Hill function was fitted to the fold change of mean gene expression compared to dose 0. Optimization was performed using "optimx()" function of the R package optim, selecting for the best optimization algorithm by the smallest residual sum of squares. Hill coefficient is shown beside the gene name. The scatter plot of normalized gene expression is shown on the right side. Red denotes mean.

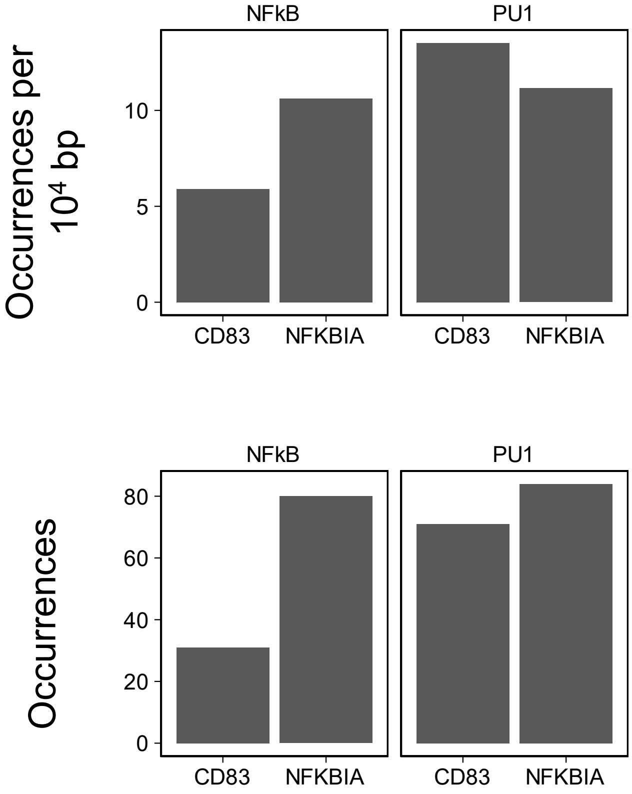


**Supplementary Figure 13. Motif occurrences for SE associated with *NFKBIA* and *CD83*.** Motif number of PU.1 and NF-kB at merged SE regions for both *NFKBIA* and *CD83* calculated using Homer (See methods).

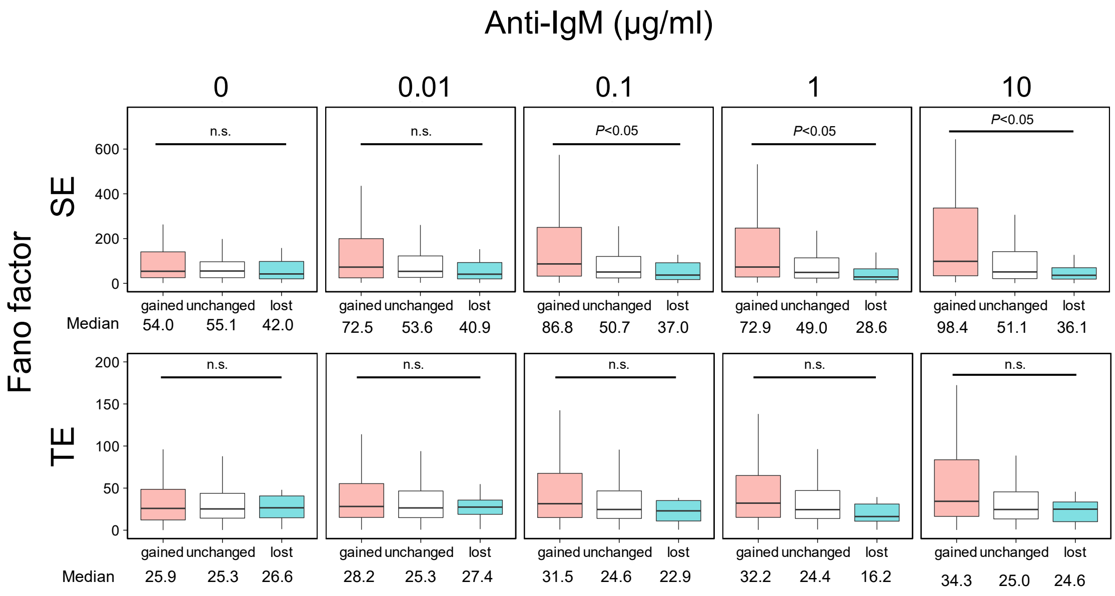


**Supplementary Figure 14. DEGs show higher Fano factor in gained SE.** Boxplot of Fano factor at all dose points for genes associated with TE and SE. Number of genes: Gained SE, 82; Unchanged SE, 100; Lost SE, 42; Gained TE, 242; Unchanged TE, 384; Lost TE, 21. *P*-value was calculated using one-way ANOVA with undersampling (n = 21), n.s.: not significant.

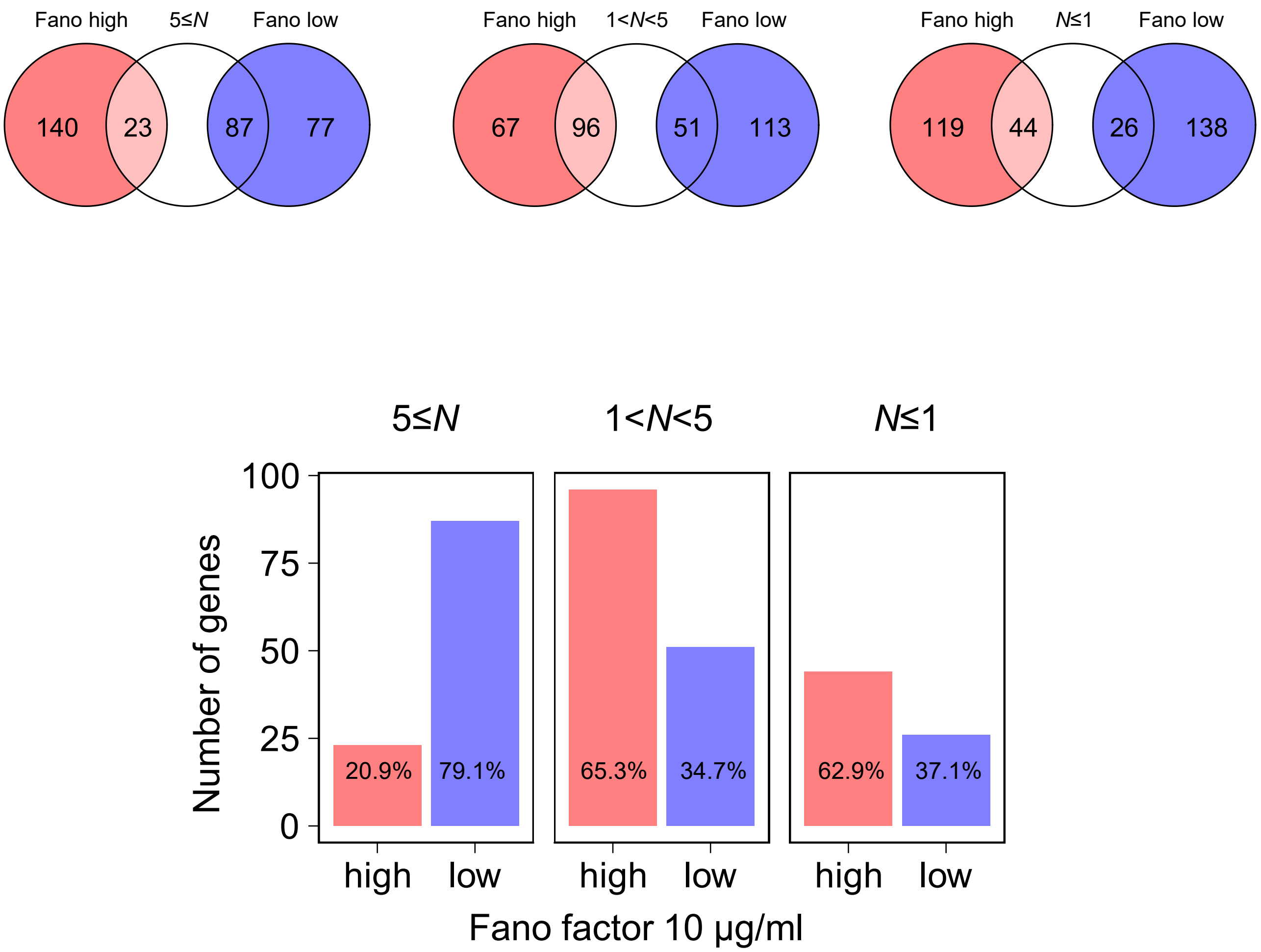


**Supplementary Figure 15. Number of genes and Hill coefficient assignment from fitting.** Venn diagram and bar plot of the number of genes with assigned Hill coefficient (*N*) below 1, above 5 and in between and subsequent Fano factor (10 µg/ml) below (low) and above median (high).

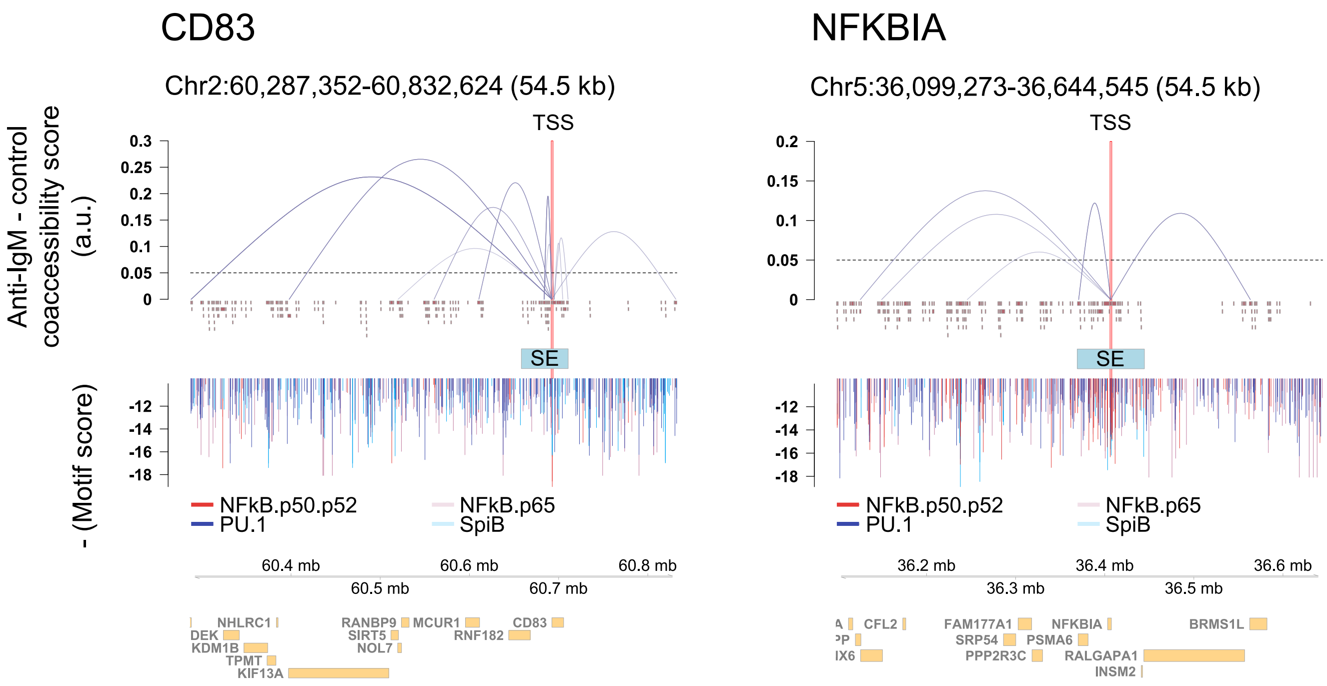


**Supplementary Figure 16. Coaccessibility of CD83 and *NFKBIA*.** Track view of *NFKBIA* and *CD83* coaccessibility +-1kb around the annotated transcription start site of and regions outside of the SE.

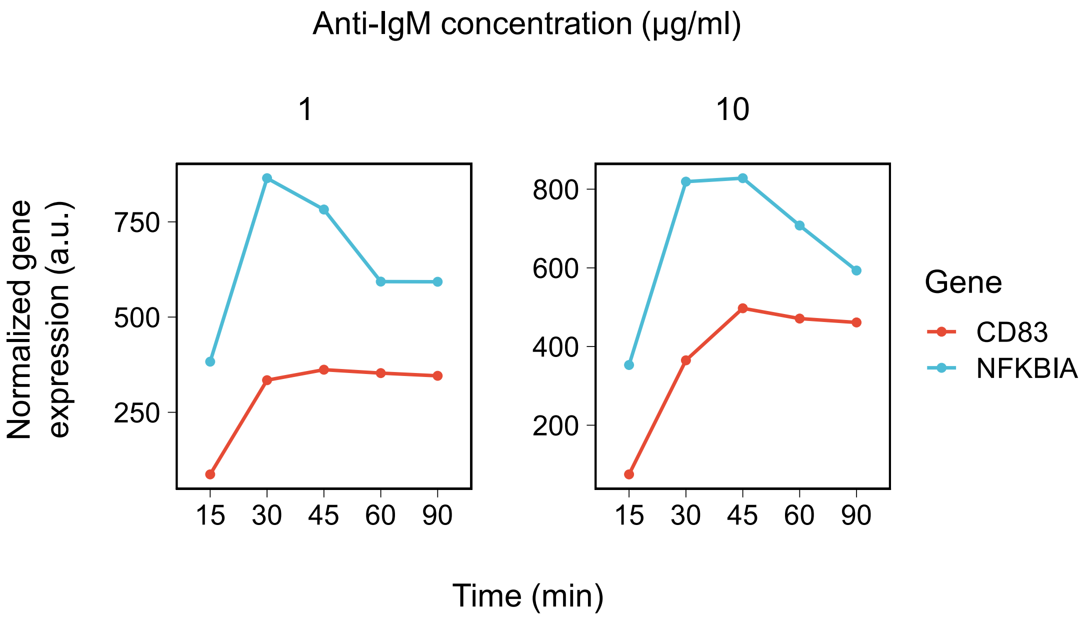


**Supplementary Figure 17. Time-course microarray analysis of *CD83* and *NFKBIA*.** Time-course normalized gene expression of *CD83* and *NFKBIA* (Chiang et al., 2020).

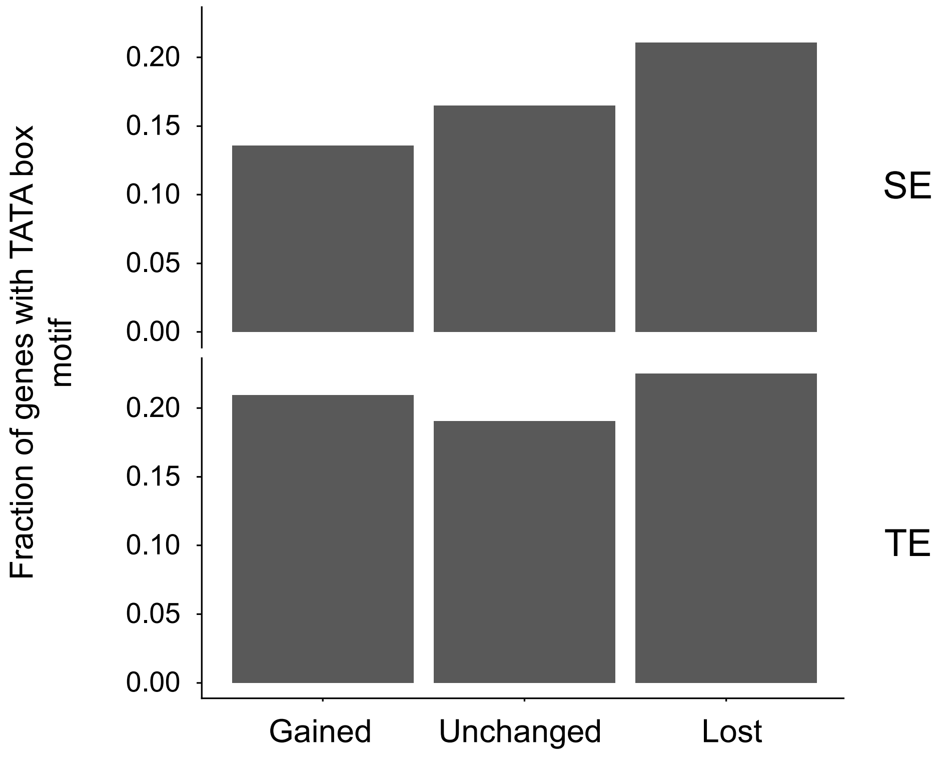


**Supplementary Figure 18. Motif occurrences of TATA box at SE and TE genes promoter.** Motif number of TATA box at promoter of TE and SE associated genes calculated using Homer (See methods).

Supplementary table 1. Number of single-cell RNA-seq data used in scRNA-seq analysis for each dose point of anti-IgM.

| Anti-IgM (μg/ml) | Number of cells |
|---|---|
| 0 | 89 |
| 0.01 | 92 |
| 0.1 | 87 |
| 1 | 92 |
| 10 | 93 |

Supplementary table 2. Number of activated and inactivated cells.

| Anti-IgM | Activated | Inactivated | % Activated | Data source |
|---|---|---|---|---|
| 0.00 | 2 | 87 | 2.25 | RNA-seq |
| 0.01 | 11 | 81 | 12.0 | RNA-seq |
| 0.10 | 24 | 63 | 27.6 | RNA-seq |
| 1.00 | 70 | 22 | 76.1 | RNA-seq |
| 10.00 | 75 | 18 | 80.6 | RNA-seq |
| 0.00 | 6 | 164 | 3.53 | Imaging |
| 0.01 | 20 | 151 | 11.7 | Imaging |
| 0.10 | 127 | 113 | 52.9 | Imaging |
| 1.00 | 220 | 36 | 85.9 | Imaging |
| 10.00 | 243 | 25 | 90.7 | Imaging |

Supplementary table 3. Primers used in mKate2-BRD4S cell line construction.

| Oligonucleotide name | Sequence (5' to 3') |
|---|---|
| attB1-mKate2 (forward) | GGGGACAAGTTTGTACAAAAAAGCAGGCTTAGCCACCATGGTGAGCGAGCTGATTA |
| attB2-*BRD4S* (reverse) | GGGGACCACTTTGTACAAGAAAGCTGGGTATTAGGCAGGACCTGTTTCGGAGTCTTCGCTGTCAGAG |
| Linker-*BRD4S* (forward) | GTGCTGGTAGTGCAGCAGGTTCTGGAGAATTTATGTCTGCGGAGAGCGGCCCTG |
| Linker-mKate2 (reverse) | CAGAACCTGCTGCACTACCAGCACTTCCTCTGTGCCCCAGTTTGCTAGGGAG |

### Supplementary table 4. List of smRNA-FISH probes (5'-3').

| ***NFKBIA* exon (Quasar 570)** | ***CD83* exon (Quasar 570)** | ***NFKBIA* intron (Quasar 670)** | ***CD83* intron (Quasar 670)** |
|---|---|---|---|
| TCATGTGCGGGTTTCGAAAG | AAATGTAAGGCTCCCTACTT | TGTTAATGCCAGACTCCTAA | GATTATTTGCCATATGTGGC |
| AAAAAAAGGGAAGCGCCCCA | CCTACATAATCCTTTCTGTC | GATACTCCTATCTACTCTGG | TACCATGCCCCAAGAAAATT |
| AAACCTCTCCTCGGGAATTC | AGGCTGCACATGGAACAATC | CAAATTCTACATGCCCTAGC | CTTACAACCATTCTTCTGGT |
| CCTAACATCCATCCAACTAC | ACCAAGAGACAGCATCACTG | CAGTATGTCTGCAACTTCTA | CAGTGTAGACATCTACACCA |
| AAGAGGCTGCAGTAGGACAA | AAAATCAGCTCCTTTCTGGG | CTGAAATGGTTACATGTGCT | CAGAGAAGCTGCTGAGTGAT |
| CCATTGCATCGCGTATGAAA | TGTGACTTAGTTTGCAGCTA | ATGGCACTAAGACCATCACA | CCCATCATCACTTCAGAATT |
| CATCCCAATGGAGAGGAATG | GCAGCACACACTATGAACTC | ACCAAAGCTGGGAACACCAA | CTAATGCTGCAGTTTAGCTT |
| TCCATATGAAGGGTATCATT | GCATGGCAAAACCTTTCTTT | CCACCAATTTATTTTATCTC | GCTTGTTTCGTAGAGTTTGA |
| CCTCCAGATATCAGACATTC | CTAGTGTTGCACTGCAATTC | TTCTGTAGGTATCAGCTCAG | GTGTTCTCATGTGTTTGCAA |
| TCTCTAGTCTGGAAGCTACG | AATGCTTTTGGCTTTTAGCT | AATCACAGCGTGAAGCTACC | TCAAATCTTCATCTGGTGGC |
| CAACGGCAAGTACTGCAAGT | GCAAAAATGCATCAGGTCCT | TTCCCCAGTGTATTATTTAC | AGCACTGTAAAGGTGGTCAC |
| CAGTGGCTTTTGAGGACTTC | AGCAAATCACTTTTTCCCAC | TTTACTCTTAAGCTCATTCT | AGCCCAAAAGGACAAGTCTT |
| ACGTCCAAATACACAAACCC | TCTTAATTAGCCCTTTGAGG | TTATATCCTTGCCCTAACTG | CCTAAATTCAGCTTCTCTCA |
| GAGTTATGTGACAAGCGTGG | AGATTTAGCCTAACAGCTGT | TTTGATGGTTGGCAGCGCAA | AACACCGAAAGCTGGTGTTC |
| AAGGCAGGAGGGAATGCACA | TACATGGTCGAAGCATGCAG | TTACTTCATGGGCTGTTATC | CCACAGCTGACTCAATACAT |
| GATACTTCACTCGAGGGCTC | CCTTTTTACATTCAGCTTCA | AGTTACATAGGGAGGACGTC | ATAGATGGTCTGTTTCTTCC |
| CTCAGCACTGCAATGGGAAC | CAAACCCATGCCTACAAACA | TGCAGCACCAGTCAAACAAA | GAGGCTTCATTTCATGTAGT |
| AAGTCAGGTACTGCATTGCT | CCACAGAAATTCAAGCCAGT | CAGGATGTGACTCTTCTTTT | TAGAAGCCTCATGTGGATGA |
| CTGAACCCACGACATCTTAA | ATGACTTCCTCCTCTAATTT | ATGCCCCATAACTTGTTTAA | CCTGCAGGAAATGTCTGATG |
| ATGACCATGAGCAAGGAGGC | GTGACACGTCTGTATGATCT | TGATAGGACAGGGCACTATC | CCACTATGTAAGTGCATCTC |
| AATTCAAAGCCCACGAGAGC | TCAGTCATGAATGCGTTGGA | AGTCTGTAAACCAACCAGGG | CTTCTCTTGCAGAGATAGGA |
| ACTGCTATGAGTACTGCAGA | AGTCAGAAAAGTCCCTGACT | CAAAACAGCTTTCCAGCTTC | CTTCCTATGTTTCAGCTTAT |
| ACAACTTCCTCCTCCAAAAA | GCATTTTGCAGCACTAAAGC | GTATGGATAACTGGGTTGGA | GATACGATTCATCCAGGACA |
| TTCCAGTGTTTAACTATGCC | ATCTAGAATCCTGGCAATGC | TTCCAGTGTTTAACTATGCC | GGCTCTTCATCATACTGATT |
| GGAAAGATTTCTTAGGGGGC | TACTCTATATCCAGCCAGAA | ACAACTTCCTCCTCCAAAAA | CTGACAAGCCTAAACTCCAG |
| GTATGGATAACTGGGTTGGA | CCTTTTCCCATAAAGGCAAA | ACTGCTATGAGTACTGCAGA | AGTCTTCACATCACGAGAGT |
| CAAAACAGCTTTCCAGCTTC | ATTTCTGGTTTCTGACTGGA | AATTCAAAGCCCACGAGAGC | CTGCCTGGTGGAGTGAAAAA |
| AGTCTGTAAACCAACCAGGG | CCTCACATGTATTCTGTGAT | ATGACCATGAGCAAGGAGGC | TCAGCTTCTTAACAGCAGTT |
| TGATAGGACAGGGCACTATC | GCTTCAGTGAAACTTATGCC | CTGAACCCACGACATCTTAA | CATTTGTTAGCCAACTAGCA |
| ATGCCCCATAACTTGTTTAA | TTTTCCTTTAAGGGTGGTTG | AAGTCAGGTACTGCATTGCT | GTAGTGCATAATGGTGCTTT |
| CAGGATGTGACTCTTCTTTT | TAGAACCCCATGAAGTTTGG | CTCAGCACTGCAATGGGAAC | AAGCACATCACTGACAGGTT |
| TGCAGCACCAGTCAAACAAA | GGTGAGCTGTACACTGTAAC | GATACTTCACTCGAGGGCTC | CACTTCTTCCTTCTTCATAT |
| AGTTACATAGGGAGGACGTC | TCTGCTGCATAAAACTCTCC | AAGGCAGGAGGGAATGCACA | TTGTTCACTGATGAGCATGC |
| TTACTTCATGGGCTGTTATC | GCTGTTAGTGTTGGAAATCC | GAGTTATGTGACAAGCGTGG | CTGTTTAAAACCCGGAGTGT |
| TTTGATGGTTGGCAGCGCAA | TTGGTACAGTCATCAGTCTG | ACGTCCAAATACACAAACCC | TCATGTGCGTTGATGAGAGT |
| TTATATCCTTGCCCTAACTG | ACCTCACCAAAACACACCAA | CAGTGGCTTTTGAGGACTTC | CGTTTTGAATCTCTTCTGCT |
| TTTACTCTTAAGCTCATTCT | GATGCCCAAACCAAAGCAAT | CAACGGCAAGTACTGCAAGT | CTGGGAGACATACTCTCTTT |
| TTCCCCAGTGTATTATTTAC | TGCGGGTGCAATTCTAAGAT | TCTCTAGTCTGGAAGCTACG | AAAATCCCGAGGCAAGTCAG |
| AATCACAGCGTGAAGCTACC | AAATACTGTGACAGCATCCC | CCTCCAGATATCAGACATTC | TTCATCTTCTATTCTAGGGC |
| TTCTGTAGGTATCAGCTCAG | ATCCTATGAGTCAACTCTCT | TCCATATGAAGGGTATCATT | TTCCTGATCCTCAGTTGAAA |
| CCACCAATTTATTTTATCTC | TGGTTACACGTGGGTCAATA | CATCCCAATGGAGAGGAATG | CAAGTCCTTTTGGATGACGA |
| ACCAAAGCTGGGAACACCAA | CTATGATGGCTTTTAGGCTG | CCATTGCATCGCGTATGAAA | TCAAGGACTTTCCATGCTAT |
| ATGGCACTAAGACCATCACA | ACTTTACCCAACAAGAGGGA | AAGAGGCTGCAGTAGGACAA | ACAGGACAGCAAAGCTTCTT |
| CTGAAATGGTTACATGTGCT | TTTTTTCTGTTATTGGGGGG | CCTAACATCCATCCAACTAC | CATGTCACAGCAACATCTGG |
| CAGTATGTCTGCAACTTCTA | CCTGTGTTTGCATGAGGAAA | AAACCTCTCCTCGGGAATTC | CAAGCTCCAAACATTGCACA |
| CAAATTCTACATGCCCTAGC | ATCTGAGTGCAACGGTTACA | AAAAAAAGGGAAGCGCCCCA | TAGAGTGTAGGCTGCTGAAG |
| GATACTCCTATCTACTCTGG | GTGCTTACATTCTATCGGTA | TCATGTGCGGGTTTCGAAAG | GGGTATTGCTGCTTCAAATC |
| TGTTAATGCCAGACTCCTAA | TTAGTACATGTCCATTCCTA | | |

Supplementary table 5. Primers used in qPCR.

| **Oligonucleotide name** | **Sequence (5' to 3')** |
|---|---|
| *CD83* (forward) | ACCCTGTGCAATGTTTGGAG |
| *CD83* (reverse) | CTGGTAGGCGATCGAGGAAT |
| *NFKBIA* (forward) | TTCACGAGGAAAAGGCCCTG |
| *NFKBIA* (reverse) | TCTGGCTGAGGTTGTTCTGG |
| *GAPDH* (forward) | AGGTGCTGAGTATGTTGTGGAGTC |
| *GAPDH* (reverse) | GTGGTGCACGATGCATTGCTGACAAT |
